# Epithelial Stem Cell Fate Determines Chemoradiotherapy Response in Rectal Cancer

**DOI:** 10.64898/2026.07.15.736775

**Authors:** Nick Li, Fiza Ishaqwala, Thomas A. Wright, Alistair Wilkinson, Petra Vlckova, Katherine Trevers, Rhianna O’Sullivan, Shauna Crampsie, Ewa Basiarz, Sierra Vanderkamp, Ashley K. McCulloch, Aurélie Dobric, Smita Krishnaswamy, Bart Vanhaesebroeck, Glasgow Serial Sampling Consortium, Campbell S. D. Roxburgh, Maria Hawkins, Christopher J. Tape

## Abstract

Rectal cancers are often treated with neoadjuvant chemoradiotherapy (CRT), yet 85% of patients do not achieve a pathological complete response. To identify the molecular determinants of CRT response, we profiled the single-cell signalling, DNA-damage, cell-cycle, apoptotic, and cell-fate responses of 2,769 patient-derived organoid cultures treated with CRT, cancer-associated fibroblasts (CAFs), and signal-rewiring agents. We find that CRT response is determined by stem cell-fate. CRT triggers comparable DNA-damage in isogenic proliferative (proCSC) and revival (revCSC) colonic stem cells, but proCSC retain damage and die whereas revCSC resolve damage and persist. Both CRT and CAFs drive proCSC to a common treatment-resistant revCSC fate and high revCSC predicts worse survival in patients. Pharmacologically constraining stem-cell plasticity increases CRT sensitivity, and Spatial Perturbation of ARrayed Tumour Assembloids (SPARTA) confirms YAP/TEAD inhibition improves chemotherapy responses in human stromal-tumour models. These results suggest that cancer cell-fate, not genotoxic damage itself, ultimately governs response to standard-of-care chemoradiotherapy.

**HIGHLIGHTS:**

- Rectal cancer stem cell-fate determines chemoradiotherapy-induced apoptosis
- proCSCs retain DNA-damage and die, whereas revCSCs repair damage and persist
- CAFs and chemoradiotherapy converge on a common chemo-radioresistant revCSC state
- SPARTA reveals TEAD inhibition blocks DNA-repair persisters in stromal assembloids

## Introduction

Colorectal cancer (CRC) is the 2^nd^ most common cause of cancer death worldwide [1], with an alarming, rising incidence in young adults [2]. Approximately 30% of CRC tumours are rectal cancers and unlike proximal bowel tumours that can only be treated with systemic neoadjuvant therapies, rectal tumours can also be treated with localised ionising radiation. Locally advanced rectal cancer (LARC) is broadly defined as a cancer in the rectum that involves or threatens the mesorectal fascia. Standard-of-care curative treatment for LARC is neoadjuvant chemotherapy plus radiotherapy (nCRT) (comprising fractionated radiotherapy with systemic fluoropyrimidine), followed by surgery to remove the rectum and surrounding lymph nodes (total mesorectal excision) [3]. This approach results in a pathological complete response (pCR) in up to 15% of patients, but only a partial (58%) or poor (27%) response in 85% of patients [4, 5]. Whilst higher rates of pCR may be achieved using radiation with combination chemotherapy, or total neoadjuvant therapy (TNT), these regimens are heterogenous and the sensitisation of poor responders remains an obstacle [6, 7, 8]. Patients with inadequate tumour regression within the bounds of the mesorectal fascia have unfavourable outcomes, as total mesorectal excision is no longer curative in this setting [9, 10]. Consequently, understanding the molecular determinants of nCRT response is crucial for the effective treatment of LARC [11].

CRC tumours arise from oncogenic mutations in colonic epithelia that drive cell-autonomous proliferation and tumour expansion. However, recent evidence from multiple laboratories has revealed that CRC cells exhibit marked non-genetic phenotypic plasticity [12, 13, 14, 15], enabling cells to occupy a spectrum of isogenic cell-fates that include proliferative colonic stem cells (proCSC), revival colonic stem cells (revCSC; also termed regenerative [12] or onco-foetal stem cells [16]), and multiple non-canonically differentiated secretory and absorptive lineages [14, 15, 17, 18]. proCSC can express canonical colonic stem cell markers (e.g. *LGR5*^*+*^, *OLFM4*^*+*^, *EPHB2*^*+*^) but are hyper-proliferative (*RRM2*^*+*^, *MKI67*^*+*^, *TOP2A*^*+*^) relative to homeostatic epithelial stem cells due to increased cell-autonomous WNT/β-Catenin and PI3K signalling [14]. revCSC are transcriptionally similar to foetal intestinal stem cells (e.g. *ANXA1*^*+*^, *EMP1*^*+*^) and are characterised by increased YAP, TGF-β, AP-1, and NF-κB signalling [18]. Our current understanding of CRC suggests that cell-autonomous somatic oncogenic mutations (e.g. *APC, TP53*, and *PIK3CA*) push cells towards a mitotic proCSC state, whereas oncogenic *KRAS* [19, 20] and signals from cancer associated fibroblasts (CAFs) [21, 22] in the tumour microenvironment (TME) can induce revCSC. While proCSCs drive tumour growth, revCSCs are enriched at the invasive front of CRC tumours [23], where they contribute to metastasis [24] and provide a developmental gateway to non-canonical differentiation in advanced disease [15]. This non-genetic plasticity allows CRC tumours to expand when unchallenged (via proCSC) and persist under stress (via revCSC) [18].

Given the crucial role of non-genetic plasticity in CRC initiation and progression, we hypothesised that proCSC to revCSC transdetermination could determine CRT response in LARC. To test this we treated a panel of rectal cancer patient-derived organoids (PDOs) with systematic combinations of chemotherapy and radiotherapy in the presence or absence of CAFs and measured early (3 hour) and late (48 hour) responses in DNA-damage, cell-cycle activity, post-translational modification (PTM) signalling, and apoptosis at single-cell resolution. Through single-cell analysis of 2,769 PDO cultures, we find that CRT response is highly patient-specific, with some PDOs undergoing rapid DNA-damage and apoptosis, while others survive CRT. We find that the proCSC-revCSC stem cell admixture is the major determinant of CRT-induced apoptosis. proCSC experience prolonged pH2AX [S139] DNA-damage response resulting in apoptosis, whereas revCSC rapidly resolve DNA-damage and survive. We find that CAFs and CRT converge rectal cancer epithelia into a similar revCSC state – suggesting that proCSC-revCSC transdetermination is a generalised response to cell-extrinsic stress. Crucially, we show that rewiring stem cell-fates by either increasing access to proCSC (via PI3K activation) or blocking access to revCSC (via YAP/TEAD inhibition) can increase radiosensitivity. Finally, we developed a new Spatial Perturbation of AR-rayed Tumour Assembloids (SPARTA) technology platform to measure single-cell drug responses in patient-derived assembloids (PDAs) and found that YAP/TEAD inhibition blocked access to chemotherapy-induced cycling DNA-damage persister cells in a stromal tumour microenvironment. Collectively, these results demonstrate that stem cell-fate is a major determinant of LARC CRT response and reveal that pharmacologically constraining epithelial stem cell admixture can increase chemo-radiosensitivity.

## Results

### Single-cell Signalling, Cell-fate, and DNA-damage Responses of Rectal Cancer PDOs to CRT

Rectal cancer PDOs can accurately model patient-specific chemo-radiosensitivity [25, 26], but previous studies have not defined the molecular determinants of CRT response. To define patient-specific responses to CRT, we first treated 10 MSS rectal cancer PDOs (Table 1) with systematic combinations of chemotherapy (5-FU, 5-FU + oxaliplatin (FOLFOX), 5-FU + SN-38 (FOLFIRI), 5-FU + SN-38 + oxaliplatin (FOLFOXIRI)) and 16 Gy irradiation (IR). To investigate the role of key proCSC and revCSC signalling pathways, PDOs were cultured +/-a PI3K activator [27] or +/-a YAP/TEAD inhibitor [28] respectively. Finally, all PDOs were cultured +/-CRC CAFs (+/-anti-fibrotic kinase inhibitor Nintedanib) to study the effect of intercellular stromal signalling on CRT response. To measure both acute and downstream treatment responses, all PDOs were assessed at 3 hours and 48 hours following 0 Gy or 16 Gy IR in triplicate (Figure 1A). All PDOs cultures were analysed using thiol-organoid barcoding *in situ* mass cytometry (TOB*is* MC) [29, 30] to resolve DNA-damage, cell-cycle, PTM signalling, stem cell-fate, and apoptosis at single-cell resolution (2,400 perturbations total, 3,600 cell-type-specific (epithelial or CAF) datasets, 12,096,814 single cells) (Figure 1B) (Table 2). To measure platinum-based chemotherapy responses (e.g. FOLFOX) by MC, we developed a new TOB*is* 2.0 barcoding system exchanging ^196^Pt-Cisplatin and ^198^Pt-Cisplatin for mDOTA-^103^Rh and BABE-^108^Pd. Finally, high-dimensional single-cell treatment effects were computed using Trellis, a rapid and scalable tree-based optimal transport system [21] (Figure 1C).

**Table 1.** PDO mutations and clinical metadata.

| Patient | Mutated Genes | MSI-H | Stage | Location | Cell Model Passport |
| --- | --- | --- | --- | --- | --- |
| PDO 05 | *APC, KRAS, PIK3CA, B2M, TCF7L2, PDGFRA* | No | IIIC | Rectum | HCM-SANG-0266-C20 |
| PDO 21 | *KRAS, SMAD4, ARID1A, PIK3R1, CTNNB1, NOTCH2* | No | IIA | Rectum | HCM-SANG-0270-C20 |
| PDO 75 | *APC, TP53, PIK3R1, PTEN, SMARCA4, ARAF, ALK* | No | I | Rectum | HCM-SANG-0271-D12 |
| CRC 12 | *APC, TP53, ARID1B, ABL1, CHD2, KMT2D* | No | IIIA | Rectum | — |
| CRC 16 | *ARID2, BRCA2, ROS1, IGF1R, VEGFR2, PTPN11* | No | IIIC | Rectum | — |
| CRC 24 | *APC, TP53, CHD2, PTEN, NRAS, ARID1A/B, ABL1, NOTCH1* | No | I | Rectum | — |
| CRC 09 | *APC, KRAS, BRCA2, TP53, AKT1, NOTCH1, ARID1B, PIK3R1* | No | IIA | Rectum | — |
| CRC 10 | *APC, TP53, KRAS, PIK3CA, BRCA2, ABL1, ARID1A/B, CHD2, KMT2C* | No | IIA | Rectum | — |
| CRC 44 | *APC, TP53, SDHA, CHD2, PTEN, PIK3R1, ABL1, ARID1A/B* | No | IIIC | Rectum | — |
| CRC 76 | *APC, TP53, KRAS, CHD2, ABL1, ARID1B* | No | IIIA | Rectum | — |

**Table 2.** Antibody panel used in all TOB*is* MC experiments.

| Extracellular Antigen | Metal | Clone | Supplier | Antibody Registry |
| --- | --- | --- | --- | --- |
| FAP | <sup>112</sup>Cd | Polyclonal | R&D Systems | AB_2102369 |
| CD326 (EpCAM) | <sup>113</sup>In | 9C4 | BioLegend | AB_756075 |
| L1CAM | <sup>146</sup>Nd | L1-OV198.5 | BioLegend | AB_2616786 |
| TACSTD2 (TROP2) | <sup>167</sup>Er | NY18 | BioLegend | AB_2564376 |
| EPHB2 | <sup>169</sup>Tm | 2H9 | BD Biosciences | AB_2738897 |
| CHGA | <sup>170</sup>Er | C-12 | Santa Cruz | AB_2801371 |
| CD55 | <sup>171</sup>Yb | JS11 | BioLegend | AB_314859 |
| Intracellular Antigen | Metal | Clone | Supplier | Antibody Registry |
| pHistone H3 [S28] | <sup>89</sup>Y | HTA28 | BioLegend | AB_1227660 |
| Vimentin | <sup>111</sup>Cd | RV202 | BD Biosciences | AB_393716 |
| Cytokeratin 18 | <sup>114</sup>Cd | C-04 | Abcam | AB_305647 |
| Pan-CK | <sup>115</sup>In | AE1/AE3 | BioLegend | AB_2616960 |
| GFP | <sup>116</sup>Cd | 5F12.4 | eBiosciences | – |
| CK20 | <sup>141</sup>Pr | D9Z1Z | CST | AB_3101888 |
| cCaspase 3 [D175] | <sup>142</sup>Nd | D3E9 | CST | AB_10897512 |
| RRM2 | <sup>143</sup>Nd | E7Y9J | CST | AB_2140946 |
| pATM [S1981] | <sup>144</sup>Nd | D25E5 | CST | AB_2798100 |
| pNDRG1 [T346] | <sup>145</sup>Nd | D98G11 | CST | AB_10693451 |
| OPTN | <sup>147</sup>Sm | E4P8C | CST | AB_3073769 |
| CDK1 | <sup>148</sup>Nd | A17 | Thermo | AB_2533105 |
| p4E-BP1 [T37/T46] | <sup>149</sup>Sm | 236B4 | CST | AB_560835 |
| pRB [S807/S811] | <sup>150</sup>Nd | J1112-906 | BD Biosciences | AB_647295 |
| pBAD [S112] | <sup>151</sup>Eu | 40A9 | CST | AB_560884 |
| DACH1 | <sup>152</sup>Sm | 3B6D2 | Thermo | AB_2883419 |
| ANXA1 | <sup>153</sup>Eu | D5V2T | CST | AB_2799031 |
| SOX9 | <sup>154</sup>Sm | 3C10 | Abcam | AB_2493181 |
| pAKT [S473] | <sup>155</sup>Gd | M89-61 | BD Biosciences | AB_2737620 |
| pNF-κB [S529] | <sup>156</sup>Gd | K10-895.12.50 | BD Biosciences | AB_647284 |
| Isotype Control | <sup>157</sup>Gd | MOPC21 | Abcam | AB_2736846 |
| pP38 [T180/Y182] | <sup>158</sup>Gd | D3F9 | CST | AB_2139682 |
| pCHK2 [T68] | <sup>159</sup>Tb | C13C1 | CST | AB_2080501 |
| KI67 | <sup>160</sup>Gd | D3B5 | CST | AB_2687446 |
| pLATS1/2 [S909/S872] | <sup>161</sup>Dy | EPR27262-6 | Abcam | – |
| pH2AX [S139] | <sup>162</sup>Dy | D7T2V | CST | AB_3675209 |
| p21 Waf1/Cip1 | <sup>163</sup>Dy | 12D1 | CST | AB_3099451 |
| TOP2A | <sup>164</sup>Dy | D10G9 | CST | AB_2797871 |
| pS6 [S240/S244] | <sup>165</sup>Ho | D68F8 | CST | AB_10694233 |
| pGSK-3β [S9] | <sup>166</sup>Er | D85E12 | CST | AB_10013750 |
| pSMAD2/3 | <sup>168</sup>Er | D27F4 | CST | AB_2631089 |
| BIRC3 | <sup>172</sup>Yb | OTI3G4 | Thermo | – |
| pP53 [S15] | <sup>173</sup>Yb | 16G8 | CST | AB_331741 |
| cPARP [D214] | <sup>174</sup>Yb | D64E10 | CST | AB_10699459 |
| pCHK1 [S345] | <sup>175</sup>Yb | 133D3 | CST | AB_331212 |
| Cyclin B1 | <sup>176</sup>Yb | GNS-11 | BD Biosciences | AB_395290 |
| 2meH3 [K4] | <sup>209</sup>Bi | C64G9 | CST | AB_10205451 |

**Figure 1.**
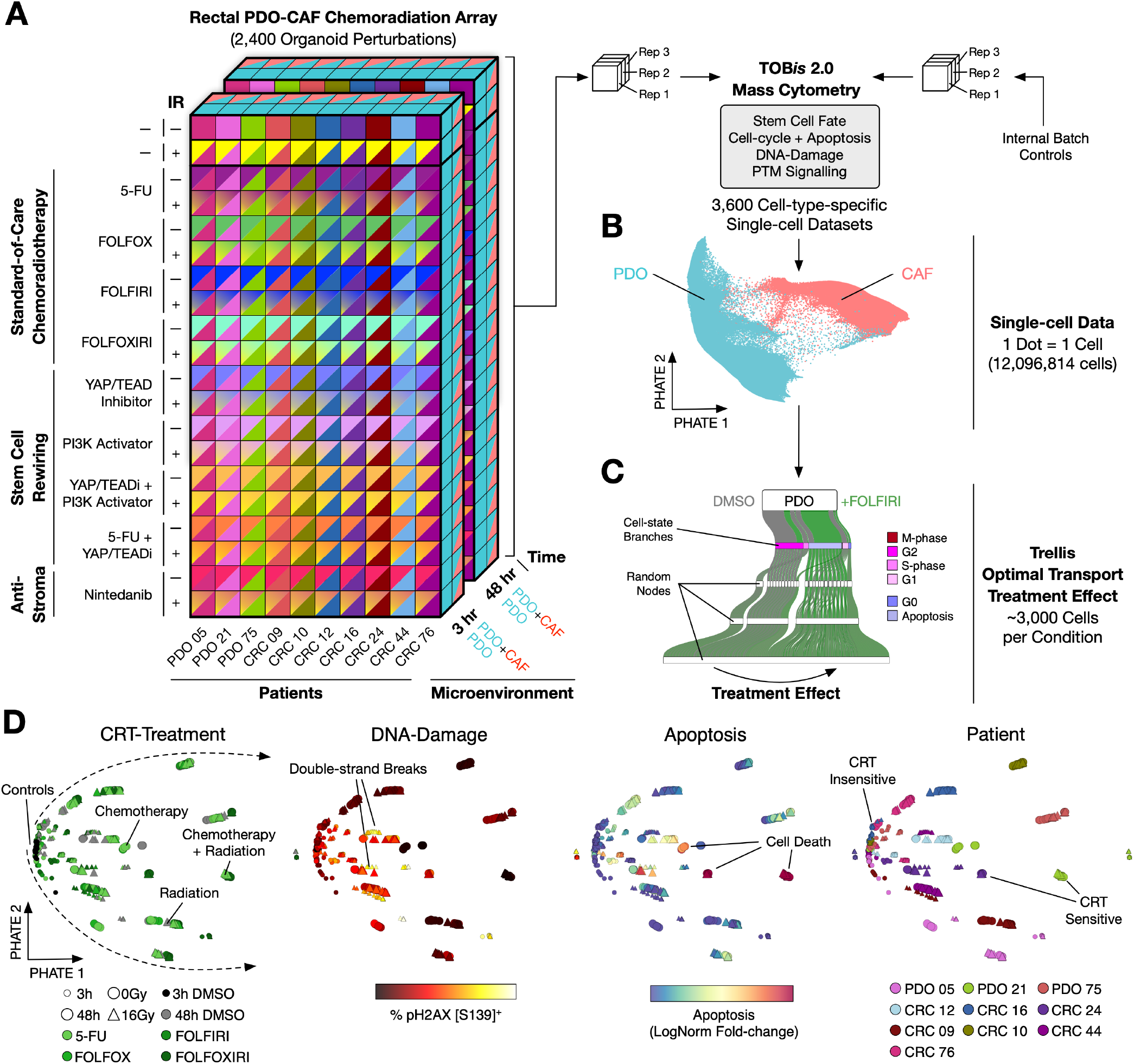
Single-cell Signalling Analysis Reveals Highly Patient-specific Rectal Cancer CRT Responses. **A)** Experimental overview. 10 rectal cancer PDOs were treated with irradiation (IR) +/-5-FU, 5-FU + oxaliplatin (FOLFOX), 5-FU + SN-38 (FOLFIRI), 5-FU + SN-38 + oxaliplatin (FOLFOXIRI) and analysed at 3 and 48 hours in triplicate using single-cell TOB*is* Mass Cytometry [29, 30]. **B)** PHATE-embedding of all 12,096,814 cells acquired (excluding internal controls). **C)** Sankey diagram showing how treatment effects were computed using Trellis optimal transport [21]. **D)** Trellis-PHATE embeddings of 600 experimental conditions coloured by treatment, patient, DNA-damage (pH2AX [S139]) and apoptosis (cCaspase [D175] and cPARP [D214]). 1 dot = 1 condition.

To understand how CRT affects rectal cancer epithelia alone, we first analysed 600 PDO monocultures treated with 5-FU, FOLFOX, FOLFIRI, FOLFOXIRI, +/-IR. Single-cell signalling analysis revealed that CRT response is highly patient-specific, with equivalent doses of chemotherapy and IR producing distinct levels of DNA-damage (pH2AX [S139]) and apoptosis (cCaspase [D175]^+^ and cPARP [D214]^+^) per patient (Figure 1D). These results are consistent with the recent paradigm shift suggesting that radiation dose as a sole parameter is insufficient to explain radiotherapy efficacy [31]. To quantify how changes in PTM signalling, DNA-damage, and phenotypic markers translate to therapy-induced apoptosis, we compared the Trellis *L*^1^ distance of each treatment against apoptosis for all PDOs. This analysis revealed three treatment response groups: high treatment response with high apoptosis, high treatment response with low apoptosis, and low treatment response with low apoptosis (Figure S1). These results demonstrate that PDOs experience variable levels of on-target DNA-damage, distinct signalling response, and apoptosis when treated with CRT – mimicking variable CRT responses observed in LARC patients [4, 5].

### Stem Cell Fate Determines CRT Treatment Response

To mechanistically explore why some PDOs enter apoptosis following CRT, whereas others survive, we first quantified pH2AX [S139] at single-cell resolution for all PDOs +/-16 Gy IR. We found that 3 hours after IR, pH2AX [S139] tightly correlates with the cell-cycle activity of each PDO (via % pRB [S807/S811]) (Figure S2A). We observe pH2AX [S139] in almost all actively cycling, pRB [S807/S811]^+^ cells, but in less than half of non-cycling cells (Figure S2B). However, pH2AX [S139] is a modest predictor of CRT-induced apoptosis (Figure 2A) – suggesting DNA-damage alone is not an accurate marker for chemotherapy efficacy. By contrast, both chemotherapy and radiation-induced apoptosis strongly correlated with the baseline stem cell index (SCI) of each PDO for both FOLFOX and IR (Figure 2B). SCI (first introduced by Gil Vasquez *et al*., [12]) represents the relative proportion of proCSC to revCSC cells within a CRC admixture, with SCI^High^ indicating an enrichment of proCSC and SCI^Low^ representing an increase in revCSC. While SCI^High^ PDOs (proCSC dominant) are fast-cycling, with a higher proportion of cells in S-phase (Figure S2C,D), SCI correlates poorly with other cell-cycle markers such as pRB [S807/S811], Cyclin B1, and pHH3 [S28]. This suggests that SCI is not purely driven by cell-cycle activity and is a more general characterisation of cell-fate identity (Figure S2D).

**Figure 2.**
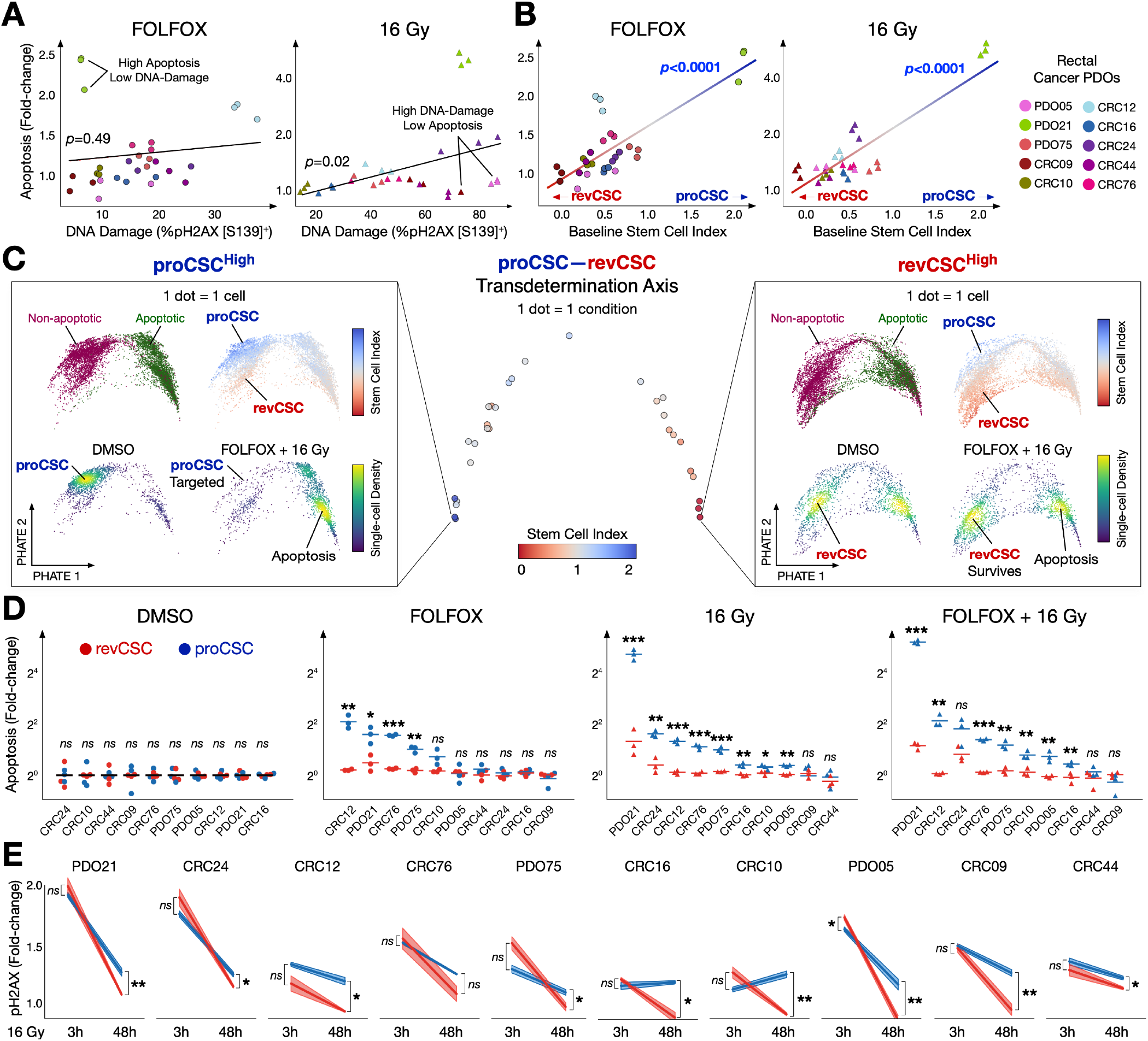
Stem Cell Admixture, Not DNA-damage Burden, Determines Rectal Cancer CRT Response. **A)** DNA-damage (% pH2AX [S139]) vs. apoptosis (fold-change) of all PDOs following FOLFOX or IR. **B)** Baseline stem cell index (SCI) vs. apoptosis (fold-change) of all PDOs following FOLFOX or IR. **C)** Single-cell PHATE of all 10 PDOs coloured by SCI and single-cell density plots of revCSC and proCSC gated cells treated with DMSO, FOLFOX, IR, and FOLFOX + IR. **D)** Apoptosis (fold-change) of proCSC and revCSC for each PDO following DMSO, FOLFOX, IR, and FOLFOX + IR. **E)** DNA-damage (% pH2AX [S139] fold-change) following IR for proCSC and revCSC cells at 3 hours (initial response) and 48 hours (late response). Paired t-tests were performed and false discovery rate correction for multiple comparisons was applied. Adjusted p-value *: p<0.05, **: p<0.01, ***: p<0.001, *ns*: non-significant.

By simultaneously measuring proCSC and revCSC markers alongside DNA-damage and apoptosis, we found that baseline SCI strongly correlates with both FOLFOX and 16 Gy IR-induced apoptosis. proCSC^High^ PDOs are disproportionately targeted by CRT, whereas revCSC^High^ PDOs resist apoptosis after CRT. To further explore the therapeutic importance of stem cell admixture, we integrated single-cell data from all PDOs, gated cells into revCSC^High^ and proCSC^High^ populations (Figure S2C), and measured their stem cell-specific responses to CRT (Figure 2C, Figure S2E). This analysis revealed that proCSC undergo a dramatic phenotypic shift towards apoptosis following CRT, whereas revCSC cells are able to maintain key survival signalling pathways under therapy (pP38-MAPK, pS6, pAKT, pNF-κB, and p4E-BP1) (Figure S2F). proCSC consistently experience more CRT-induced apoptosis than revCSC (Figure 2D). Interestingly, we find that both proCSC and revCSC display equal levels of on-target DNA-damage (pH2AX [S139]) at 3 hours post-irradiation, but proCSC cells retain significantly higher DNA-damage 48 hours later in 9/10 PDOs (Figure 2E). Collectively, these results demonstrate that within an isogenic stem cell admixture, CRT can trigger equal DNA-damage in both proCSC and revCSC, but only proCSC enter apoptosis while revCSC survive therapy. As a result, we conclude that stem cell-fate is a major determinant of CRT treatment response in rectal cancer.

### CRT Induces Cell-fate Specific DNA-damage Responses and Polarises LARC Epithelia to revCSC

Single-cell analysis of rectal cancer PDOs revealed that CRT triggers apoptosis in proCSCs, whereas revCSC survive CRT. We therefore hypothesised that CRT itself could induce stem cell plasticity. Using a time-adjusted SCI (Δ*SCI*_*adj*_) to correct for natural SCI drift during unperturbed organoid culture, we found that 5-FU, FOL-FOX, FOLFIRI, and IR therapies can polarise LARC cells towards a revCSC-dominant admixture (Figure 3A). While revCSC polarisation could in theory be caused by either cell fate-switching and/or proCSC apoptosis, we find that epithelial apoptosis did not correlate with drop in (Δ*SCI*_*adj*_) across the cohort, suggesting that CRT induces proCSC to revCSC transdetermination (Figure S3A,B).

**Figure 3.**
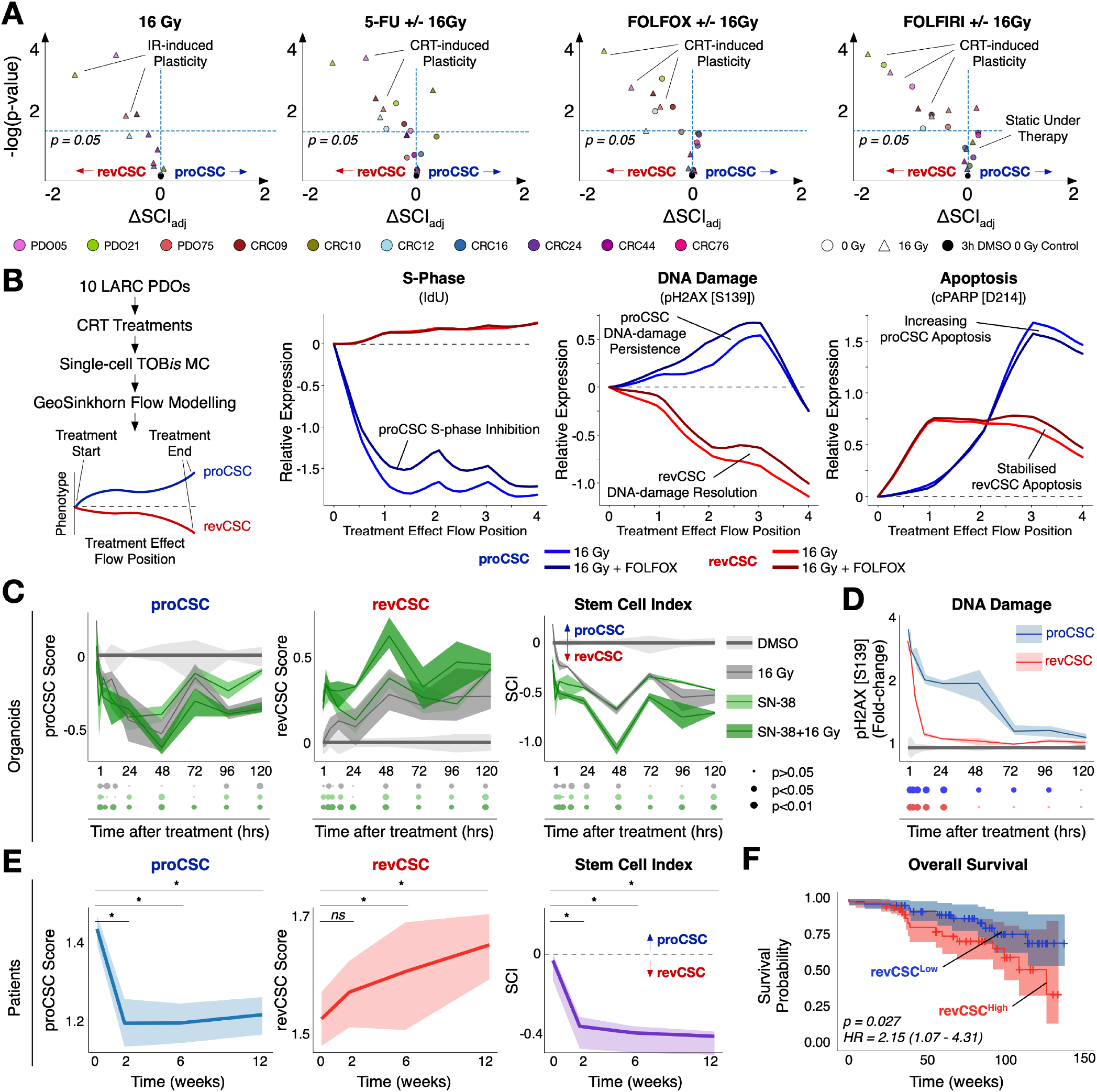
CRT Induces revCSC in Patients and Organoids, and High revCSC Predicts Worse Survival. **A)** Change in stem cell index (Δ*SCI*_*adj*_) following IR, 5-FU (+/-IR), FOLFOX (+/-IR), and FOLFIRI (+/-IR) for 10 LARC PDOs. **B)** GeoSinkhorn Flow dynamic flow modelling of S-phase, DNA-damage, and apoptosis for proCSC and revCSC cells from 10 LARC PDOs treated with IR and IR + FOLFOX analysed by TOB*is* MC. **C)** TOB*is* MC proCSC and revCSC time-course analysis of PDO21 treated with DMSO, IR, SN-38, and IR + SN-38. P-values were generated using paired t-tests of treatment condition vs control with false discovery rate correction for multiple comparisons.**D)** pH2AX [S139] fold-change after IR relative to DMSO control in proCSC and revCSC. **E)** Time course of changes in proCSC and revCSC signatures of a clinical standard-of-care rectal cancer cohort undergoing nCRT (weeks 0-6) over 12 weeks (*n* = 48). Pairwise comparisons performed with Wilcoxon signed-rank test. p-value *: p<0.05, *ns*: non-significant. **F)** Overall patient survival (S:CORT data) after nCRT for rectal cancer relative to revCSC expression (optimal cutpoint: NES 0.98, revCSC^High^ (*n*=107), revCSC^Low^ (*n*=116)).

We found that rectal cancer cells with high mitotic signalling (e.g. p4E-BP1 [T37/T46], pP38-MAPK [T180/Y182], pNDRG1 [T346]) and low Hippo signalling (pLATS1/2 [S909/S872]) that also express proCSC markers EPHB2, TOP2A, and SOX9 can transition to revCSC following CRT (Figure S3C). However, epithelial cells with increased pBAD [S112] and pP53 [S15] signalling that express revCSC-like markers ANXA1, CD55, BIRC3, and OPTN are phenotypically static under therapy. These results suggest the initial signalling state of LARC cells determines not only whether cells die following CRT, but their ability to further retreat to drug-tolerant persister-like states under therapy.

Single-cell CRT perturbation analysis suggested that proCSC and revCSC have distinct cell-fate specific responses to CRT. To explore the dynamics of cell-fate specific phenotypic response to CRT, we used GeoSinkhorn Flow dynamic flow modelling [32] to computationally model proCSC- and revCSC-specific CRT responses across 10 PDOs (Figure 3B). GeoSinkhorn Flow modelling suggested that following CRT, proCSC rapidly decrease S-phase, accumulate a high-level of DNA-damage during treatment which then drops as cells enter CRT-induced apoptosis. By contrast, flow modelling suggested that revCSC reduce CRT-induced DNA-damage quickly after treatment and have a modest apoptotic response following CRT. These results suggest that across all patients tested, proCSC and revCSC have fundamentally distinct responses to the same CRT treatment.

To empirically understand the dynamics of CRT-induced plasticity, we performed a single-cell time-course analysis of rectal cancer PDOs over 120 hours following treatment (9 time-points, 369 PDO cultures, 1,319,236 cells) (Figure 3C). Time-course analysis revealed CRT acutely reduces proCSC in the first 48 hours, followed by a gradual recovery. By contrast, CRT stably increases revCSC, resulting in an overall decrease in collective SCI during CRT treatment. In agreement with computational flow modelling, we found that while both proCSC and revCSC experience equal pH2AX [S139] immediately after treatment, revCSC rapidly resolve DNA-damage, whereas proCSCs retain high-levels of pH2AX [S139] throughout treatment (Figure 3D). These data suggest that proCSC struggle to repair CRT-induced DNA-damage and enter apoptosis, while revCSC resolve DNA-damage quickly, avoid apoptosis, and survive as drug-tolerant persister cells.

To explore whether CRT-induced plasticity is also observed in LARC patients, we quantified proCSC and revCSC gene signatures from bulk RNA-seq analysis of biopsies taken from LARC patients either before CRT, on-treatment at weeks 2 and 6, or end of treatment at week 12 (Glasgow Serial Sampling Cohort). We find that, just like PDOs, LARC patients experience a rapid drop in proCSC following CRT, accompanied by a gradual increase in revCSC (Figure 3E). As a result, nCRT induces an overall drop in SCI – enriching for chemo-radioresistant revCSC cells in LARC patients. Further analysis of the S:CORT cohort of 223 patients undergoing nCRT for LARC revealed that patients with a high revCSC signature prior to treatment experienced a worse overall survival (Figure 3F). Collectively, these results suggest that CRT eliminates proCSC cells, leading to a subsequent enrichment of chemo-radioresistant revCSC cells *in vivo*.

### Pharmacological Regulation of Stem Cell Admixture Can Sensitise Rectal Cancer Cells to CRT

Our results strongly suggest that proCSC are sensitive to CRT, whereas revCSC are chemo-radioresistant. We therefore hypothesised that either increasing access to proCSC or reducing access to revCSC could improve CRT responses. While most proliferative rectal cancer cells are pAKT [S473]^+^, pS6 [S240/244]^+^, p4E-BP1 [T37/T46]^+^, and pNDRG1 [T346]^+^ (Figure S2F), analysis of scRNA-seq data from a cohort of CRC PDOs [22, 33] revealed that putative downstream transcriptional activation [34] of the PI3K pathway significantly correlated with sensitivity to 5-FU, SN-38, and Oxaliplatin chemotherapies (Figure 4A) (Figure S4A). Moreover, in agreement with our prior work showing that PI3K activity is important for proCSC [14], PI3K-driven transcriptional activity strongly correlates with the proCSC gene expression signature (Figure 4B). We therefore attempted to enrich proCSC by activating PI3K signalling. Treating rectal cancer cells with the first-generation PI3K activator (UCL-TRO-1938) [27] resulted in increased pAKT [S473], pS6 [S240/244], and pNDRG1 [T346] signalling after 3 hours, followed by a cell-fate shift towards proCSC after 24-48 hours (Figure S4B). We next treated 10 rectal cancer PDOs +/-UCL-TRO-1938 for 24 hours before +/-16 Gy IR and measured single-cell DNA-damage, cell-cycle, PTM signalling, stem cell-fate, and apoptosis responses at 3 and 48 hours after IR using TOB*is* MC (240 conditions). Single-cell analysis revealed that UCL-TRO-1938 can increase the proportion of proCSC cells in a subset of PDOs (Figure 4C) with low baseline pNDRG1 signalling (Figure S4C). PI3K activation converted the chemoradioresistant revCSC-rich ‘cystic’ PDO05 into a ‘budding’ proCSC-rich PDO (Figure 4D) and sensitises PDO05 to IR (significant IR effect only in the presence of 1938) (Figure 4E). However, while first-generation PI3K activation can increase the proportion of proCSC cells in some PDOs, our results suggest that it does not eliminate access to the revCSC persister pool. As a result, we hypothesised that reducing access to the persister revCSC fate may provide a more comprehensive strategy for CRT-sensitisation.

**Figure 4.**
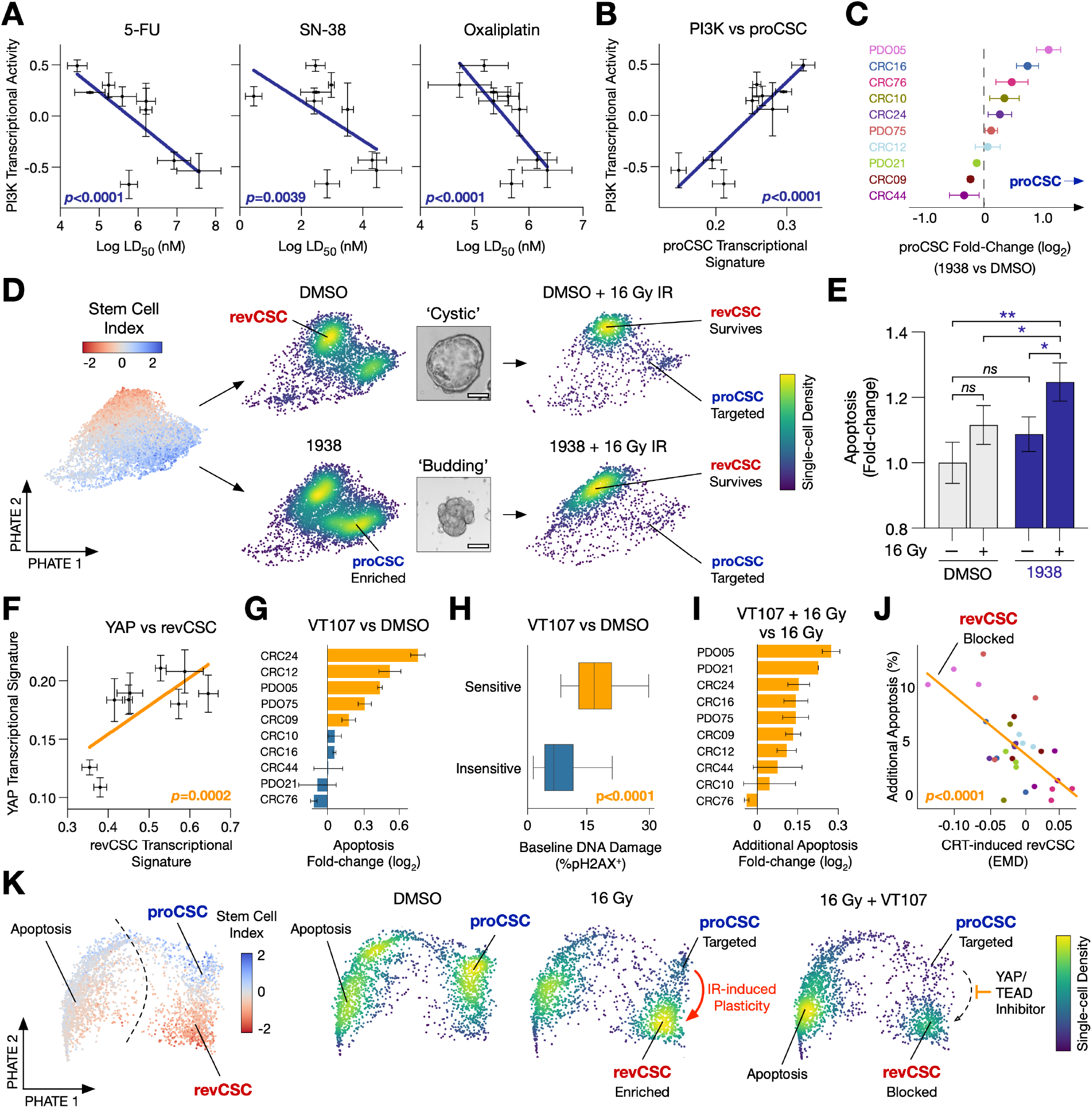
Pharmacologically Re-wiring Rectal Cancer Stem Cells to Improve CRT Response. **A)** Transcriptional PI3K activity correlated against 5-FU, SN-38, and Oxaliplatin LD_50_, and **B)** proCSC gene signature from 9 CRC PDOs (*n*=3). **C)** proCSC fold-change (log_2_) following UCL-TRO-1938 vs DMSO per rectal cancer PDO. **D)** Single-cell PHATE (coloured by stem cell index or single-cell density) and bright-field microscopy of PDO05 treated +/-UCL-TRO-1938, +/-16 Gy irradiation (IR). Scale bar = 100 µm. **E)** Apoptosis (fold-change) of PDO05 treated +/-UCL-TRO-1938, +/-IR (*n*=3). Statistical significance was assessed using t-tests with false discovery rate correction. Adjusted p-value *: p<0.05, *ns*: non-significant. **F)** Transcriptional YAP signature against revCSC signature in 9 CRC PDOs (*n*=3). **G)** Apoptosis following treatment with VT107 . **H)** Baseline DNA damage (pH2AX [S139]) in VT107-sensitive and VT107-insensitive PDOs. **I)** Additional apoptosis conferred by treatment with IR + VT107 vs IR alone. **J)** Correlation of VT107-induced revCSC expression and VT107-induced apoptosis in PDOs treated with IR. **K)** Single-cell PHATE of PDO05 coloured by SCI treated with +/-IR, +/-VT107. Linear standard curves and p-values were obtained using linear regression.

In agreement with previous studies [21], scRNA-seq data from CRC PDOs [22, 33] revealed that revCSC-dominant PDOs have increased YAP-related gene expression signatures (Figure 4F). Similarly, pLATS1/2 is increased in proCSC (Figure S2F), indicating protein-level Hippo signalling and YAP/TEAD transcriptional activity are key to the revCSC cell-fate. We therefore hypothesised that blocking access to revCSC by inhibiting YAP/TEAD signalling could improve CRT response. To test this, we treated PDOs with the TEAD palmitoylation inhibitor VT107 (Vivace Therapeutics) [28] +/-16 Gy IR and measured single-cell responses using TOB*is* MC (240 conditions). Interestingly, we find that YAP/TEAD inhibition has single-agent efficacy against PDOs with high baseline pH2AX (Figure 4G,H), suggesting a relationship between DNA-damage and Hippo signalling in rectal cancer cells. Single-cell analysis revealed that TEAD inhibition increases pS6, pBAD, pCHK2, pATM and decreases pNDRG1 and p4E-BP1 signalling (Figure S4D). Crucially, we find that YAP/TEAD inhibition increases IR-induced apoptosis in 9/10 PDOs (Figure 4I). PDOs that experience the most IR-induced apoptosis with YAP/TEAD inhibition also have reduced IR-induced access to revCSC (Figure 4J-K). Interestingly, PDOs that are not sensitive to TEAD inhibition alone still upregulate pH2AX and pCHK1 following TEAD inhibition, which then increases IR-induced apoptosis (Figure S4E). These results strongly suggest that limiting access to the drug-tolerant persister phenotypes of revCSC via YAP/TEAD inhibition can increase IR-induced apoptosis.

### CAF Radio-protect Rectal Cancer Cells by Regulating Stem Cell Admixture

Having established that chemo-radioresistant revCSC are enriched by CRT, and that access to revCSC persister phenotypes can be partially blocked by a YAP/TEAD inhibitor, we next explored whether stem cell rewiring could be effective in the presence of external signalling from the stroma. Single-cell optimal transport analysis of 2,400 PDO cultures +/-CAFs +/-CRT revealed that CAFs can induce revCSC in a patient-specific manner, and that this effect is additive on top of CRT-induced revCSC (Figure 5A). We find that CAFs can convert chemo-radiosensitive proCSC PDOs into chemo-radioresistant revCSC (Figure 5B). We find that CAF-induced epithelial plasticity is determined by the baseline epithelial stem cell index, with CAF-responsive PDOs being proCSC^High^ and poorly-responding PDOs being revCSC^High^ (Figure 5C). Interestingly, CAF-induced plasticity and CRT-induced plasticity both converge on a similar revCSC phenotype - suggesting proCSC to revCSC transdetermination is a generalised response to cell-extrinsic stress (Figure S5A,B). CAFs themselves are not altered by CRT-treatments, but are instead regulated by PDOs in a patient-specific manner (Figure S5C). Crucially, CAF-induced chemo-radioprotection strongly correlates with CAF-induced revCSC (Figure 5D/E), and not with therapy-induced DNA damage, which is unchanged by PDO+CAF co-culture (Figure S5D).

**Figure 5.**
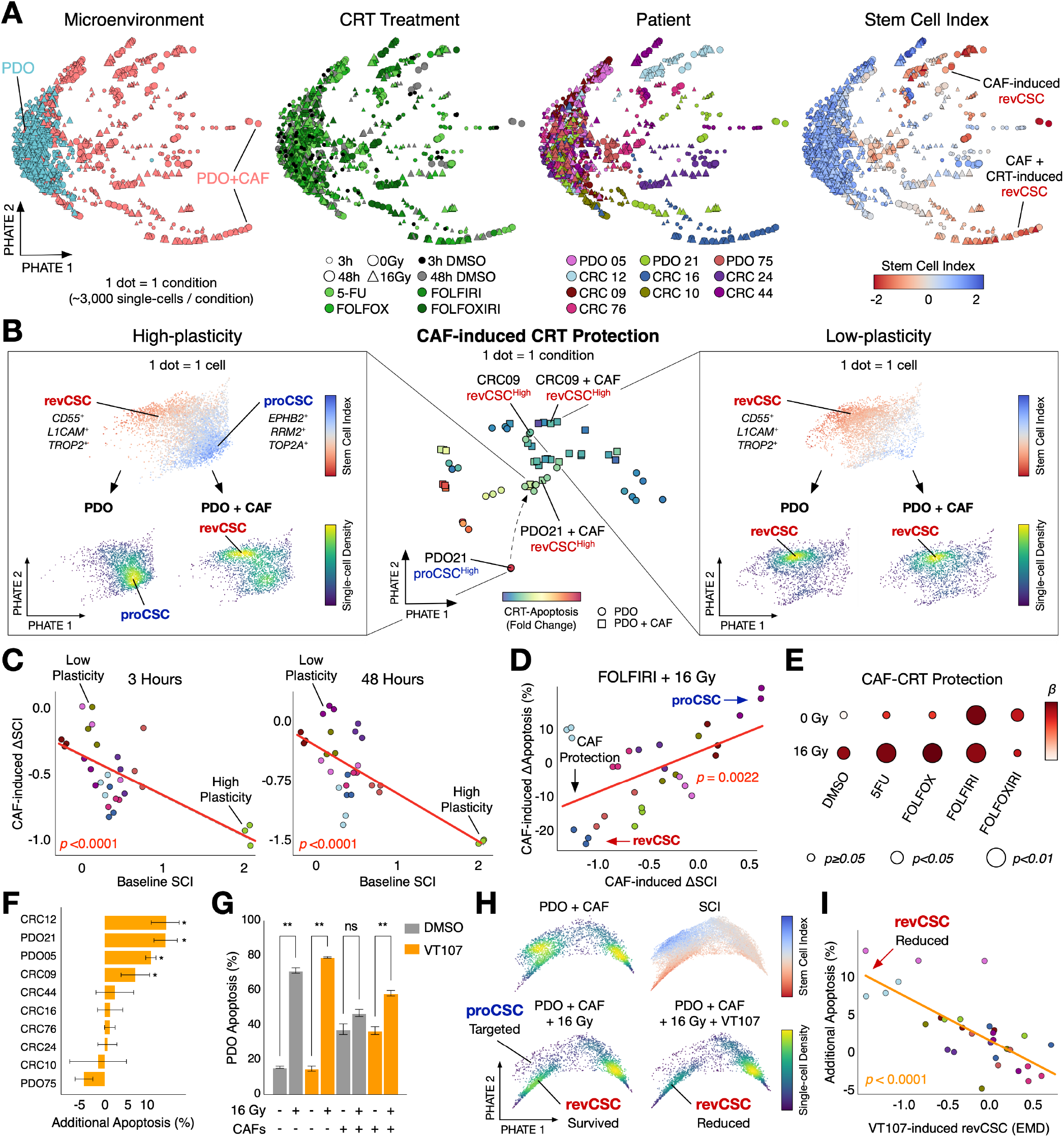
CAF-induced revCSC Drives Chemo-radioprotection and Can Be Reversed by Stromal Signalling Blockade. **A)** Paired TreEMD PHATE of 10 LARC PDOs treated with 5-FU, FOLFOX, FOLFIRI, FOLFOXIRI +/-radiation, +/-CAFs, at 3 and 48 hrs after CRT (2,400 single-cell datasets). **B)** Centre: PHATE of 10 PDOs +/-CAFs coloured by apoptosis fold-change in response to FOLFOX +/-IR. Single-cell PHATE of PDO 21 (left) and CRC 09 (right) in monoculture and co-culture with CAFs. **C)** Baseline SCI against CAF-induced plasticity. **D)** CAF-induced plasticity vs CAF-mediated protection for FOLFIRI + IR. Linear standard curves and p-values were obtained using linear regression. **E)** CAF-CRT protection effect. *β* = coefficient of linear regression. **F)** Apoptosis of PDOs in co-culture treated with +/-VT107. **G)** PDO21 apoptosis +/-CAFs, +/-IR, +/-VT107. Statistical significance was assessed using paired t-tests with false discovery rate correction for multiple comparisons. Adjusted p-value *: p< 0.05, **: p<0.01, ***: p<0.001, *ns*: non-significant. **H)** Single-cell PHATE of PDO21 in co-culture with CAFs +/-IR, +/-VT107. **I)** Correlation of VT107-induced revCSC expression (EMD of revCSC score between PDOs in co-culture +IR +/-VT107) and VT107-induced apoptosis. Standard curve and p-value obtained using linear regression.

Given the importance of CAF-induced proCSC to revCSC transdetermination in driving chemo-radioprotection, we hypothesised that blocking of CAF-PDO signalling may be able to re-sensitise PDOs. To test this, we treated PDO+CAF cultures with the anti-stromal tyrosine kinase inhibitor, Nintedanib, which is currently in clinical use as an anti-fibrosis agent for idiopathic pulmonary fibrosis [35]. CAF-induced revCSC can be partially blocked by Nintedanib, as PDOs in CAF co-culture become phenotypically more similar to their monoculture counterparts (Figure S6A) and PDOs that have been rendered resistant by CAF co-culture can be re-sensitised by Nintedanib (Figure S6B,C). Nintedanib inhibits cell proliferation and survival pathways in PDOs that are upregulated by CAF co-culture, including pAKT, pBAD, pP38-MAPK and pSMAD2/3 (Figure S6D). Interestingly, we observe an increase in DACH1 and 2meH3 [K4] with Nintedanib treatment, both features that are lost after exposure to CRT (Figure S2F). We next explored how YAP/TEAD inhibition altered epithelial chemo-radiosensitivity in the presence of CAFs. In agreement with the role of YAP/TEAD as a crucial regulator of revCSC, we found that VT107 can confer additional radiosensitivity to PDO+CAF co-cultures under CRT (Figure 5F-H). Chemo-radio re-sensitisation strongly correlates with the ability of VT107 to block revCSC (Figure 5I). Collectively, these results suggest that blocking access to CAF-induced revCSC can partially overcome chemo-radioresistance in rectal cancer.

### Spatial Perturbation of ARrayed Tumour Assembloids (SPARTA) Confirms YAP/TEAD Inhibition Blocks Drug-tolerant Persister States in Stromal Patient-derived Assembloids

CRC tumours with a high stromal content (often assigned Consensus Molecular Subtype 4 (CMS4) [36]) have poor response to radiotherapy [37] and a worse prognosis [38, 39]. While PDO + CAF co-cultures provide an excellent high-throughput model of epithelial-stromal communication, organoids lack spatial information of tumour-stroma interactions and do not model the dense fibroblast stroma found in CRC tumours *in vivo*. To overcome these limitations, we adapted murine intestinal assembloids methods [40] to develop human patient-derived assembloids (PDAs) comprising epithelial cancer cells and CAFs. PDAs self-assemble into epithelial lumens surrounded by a dense mesenchymal stroma *in vitro*, reminiscent of human CRC tumour architecture *in vivo*. To assess the molecular impact of perturbations on PDAs, we developed a custom assembloid microarray fabrication strategy to position perturbed assembloids in defined spatial positions, and then analysed assembloid microarrays with Xenium spatial transcriptomics to report single-cell gene expression for each assembloid. The resulting ‘Spatial Perturbation of ARrayed Tumour Assembloids’ technology can measure single-cell transcriptomes (5,001 genes/cell) of >100 perturbed assembloids in parallel. SPARTA therefore transforms spatial transcriptomics from a low-throughput descriptive tool into a screening-scale perturbation technology (Figure 6A).

**Figure 6.**
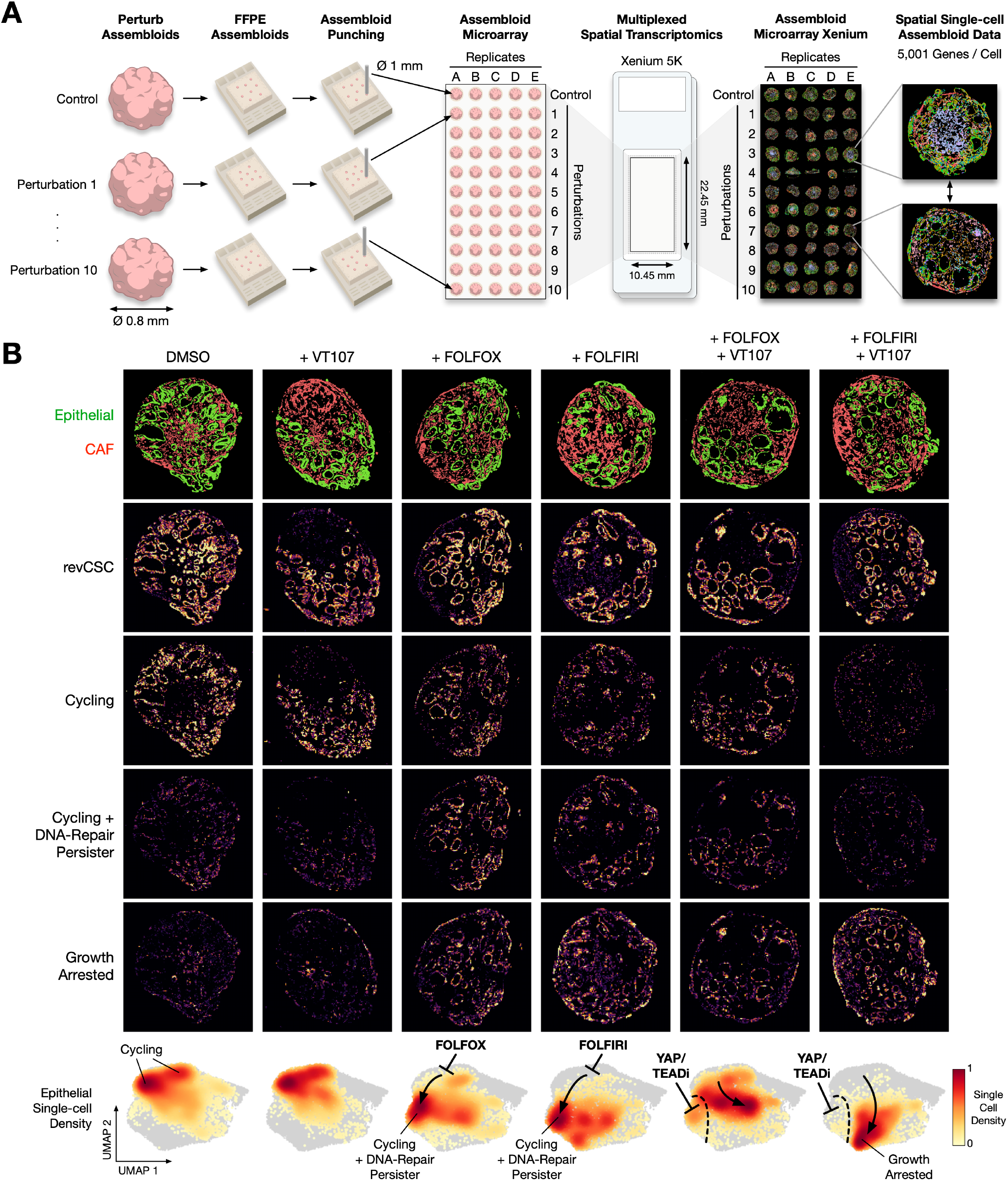
Spatial Perturbation of ARrayed Tumour Assembloids (SPARTA) Reveals YAP/TEAD Inhibition Blocks Drug-tolerant DNA-repair Persister States in Stromal Assembloids. **A)** SPARTA workflow overview. Assembloids visualised by graph-based clustering (GEX). **B)** SPARTA analysis of LARC PDAs treated with VT107, FOLFOX, and/or FOLFIRI. Images annotated with epithelial and CAF cell-types, and revCSC, cycling, cycling + DNA-repair persister, and growth arrest gene signatures. Single-cell epithelial density UMAPs for each perturbation (*n*=5).

To understand how standard-of-care chemotherapies act on highly stromal tumours we analysed rectal cancer PDAs perturbed with FOLFOX or FOLFIRI, +/-VT107 (YAP/TEAD inhibitor) with SPARTA (Figure 6B, Figure S7). SPARTA could clearly resolve epithelial and stromal cell-types and confirmed that all epithelia express a revCSC gene signature [22] (*EMP1*^+^, *L1CAM*^+^) as expected from single-cell PDO + CAF analysis (Figure 5B). SPARTA demonstrated that DMSO and VT107 treated PDAs are actively cycling (*PLK1*^+^, *CDC20*^+^, *CCNB1*^+^) with minimal evidence of cellular stress. As expected, both FOLFOX and FOLFIRI chemotherapies reduce epithelial cell-cycle activity. However, we found that FOLFOX and FOLFIRI treated PDAs do not fully exit the cell-cycle, but instead enter a cycling state with evidence of an active DNA-damage response (*MDM2*^+^, *TYMS*^+^, *FOXM1*^+^, *PCNA*^+^). This suggests that stromal-induced revCSC maintain cell-cycle activity and repair their DNA in the presence of standard-of-care chemotherapies – providing a drug-tolerant persister state. This PDA observation is consistent with the improved DNA-damage repair response profile of revCSC we observed with rectal cancer PDOs (Figure 2E, Figure 3D). Interestingly, while YAP/TEAD inhibition did not alter PDAs alone, we found that the FOLFOX- and FOLFIRI-induced repair state could be blocked by VT107. When combined with FOLFIRI, VT107 YAP/TEAD inhibition stopped cells from occupying the cycling + DNA-repair persister state and instead pushed epithelia to a highly stressed growth arrested state. These results suggest YAP/TEAD activity is important for the drug-tolerant persister properties of revCSC LARC cells in stromal tumours and pharmacological inhibition of Hippo signalling can increase chemosensitivity.

In summary, through studying the PTM signalling, DNA-damage, cell-cycle, apoptosis, and cell-fate dynamics of 2,769 rectal cancer PDOs at single-cell resolution and PDAs with single-cell spatial transcriptomics, we demonstrate that epithelial cell-fate is the major determinant of CRT response in rectal cancer. We find that proCSC experience prolonged CRT-induced DNA-damage resulting in apoptosis, whereas revCSC resolve DNA-damage faster and remain as drug-tolerant persister cells. Crucially, pharmacologically regulating stem cell plasticity via YAP/TEAD inhibition stops rectal cancer cells from accessing DNA-damage repair states, improving CRT responses. These results support a rectal cancer CRT response model whereby epithelial stem cell-fate determines CRT sensitivity and pharmacological rewiring of cell-fate plasticity can improve response to standard-of-care therapies (Figure 7).

**Figure 7.**
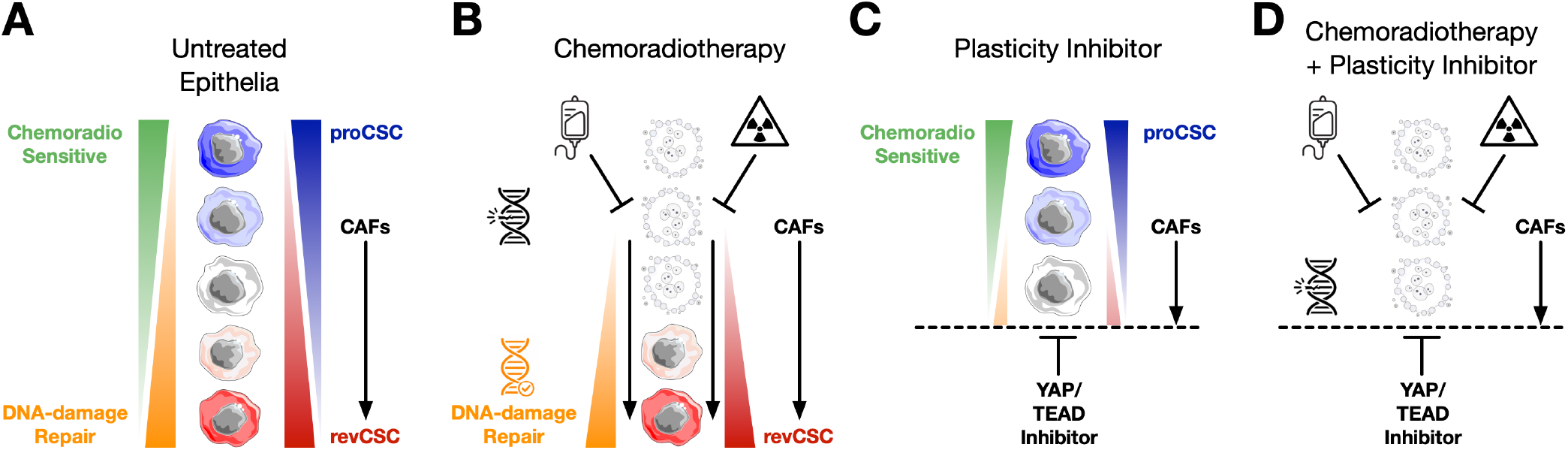
Rectal Cancer Cell-fate CRT Response Model. **A)** Untreated rectal cancer epithelia occupy a spectrum of cell-fates along the proCSC–revCSC axis. proCSC are chemoradiosensitive, damage-retaining and apoptosis-prone, whereas revCSC are chemoradioresistant, damage-resolving and persister-prone. CAF-derived intercellular signalling provides non-genetic access to revCSC. **B)** Standard-of-care chemoradiotherapy preferentially eliminates proCSC and also promotes proCSC-to-revCSC plasticity, producing a revCSC-dominant persister pool. **C)** YAP/TEAD inhibition limits access to revCSC persister phenotypes. **D)** Combining chemoradiotherapy with YAP/TEAD inhibition eliminates chemoradiosensitive proCSC while blocking therapy-induced access to revCSC DNA-repair persister states, thereby increasing therapeutic response.

## Discussion

Neoadjuvant chemoradiotherapy for LARC aims to induce DNA-damage and epithelial cell death prior to surgical resection of the primary tumour. However, clinical responses to nCRT remain unpredictable, with only 15% of patients experiencing a pathological complete response [4, 5]. Here, through systematic analysis of chemotherapy and radiotherapy responses across 2,769 organoid cultures at single-cell resolution (*>*13 million single cells), we identify epithelial stem cell-fate as a major determinant of CRT response.

In agreement with clinical observations, we find that CRT response is highly patient-specific. However, the amount of on-target CRT-induced DNA-damage did not explain therapeutic apoptosis. Instead, the baseline stem cell admixture between proCSC and revCSC determined whether LARC cells converted genotoxic damage into cell death. proCSC were exquisitely chemo-radiosensitive, whereas revCSC were chemo-radioresistant. These results suggest that current CRT regimens preferentially target the proCSC compartment of LARC tumours, leaving revCSC to survive as drug-tolerant persister cells. LGR5^+^ proCSC-like cells can occupy anywhere from 10-70% of the epithelial compartment of a CRC tumour [41] and the range of proCSC-to-revCSC within an individual’s tumour can vary substantially [12]. When combined with our results, this suggests that standard-of-care CRT can only effectively target the proCSC subpopulation of human LARC tumours.

A central implication of this work is that DNA-damage is necessary but not sufficient for CRT efficacy. Across our cohort, equivalent CRT treatments achieved different amounts of DNA-damage to different PDOs, but this variation was a poor predictor of apoptosis. More importantly, proCSC and revCSC from the same PDO line sustained equivalent early pH2AX^+^ DNA-damage following irradiation, yet only proCSC retained high-level DNA-damage and entered apoptosis. By contrast, revCSC resolved DNA-damage more quickly and survived. Thus, CRT response is not simply determined by whether a tumour cell is damaged, but by how that cell interprets and resolves the damage. Under this model, epithelial cell-fate acts as a gatekeeper between genotoxic injury and cytotoxic outcome: proCSC convert CRT-induced DNA-damage into apoptosis, whereas revCSC convert the same insult into repair and persistence.

revCSC/foetal-like epithelia have been repeatedly implicated in disease recurrence in CRC [16, 18, 42]. Our data extend this concept by providing direct evidence that proCSC and revCSC differ in their molecular response to CRT-induced DNA-damage. The higher the ratio of revCSC to proCSC within a tumour admixture, the greater the capacity of that admixture to tolerate or repair CRT-induced genotoxic stress. This provides a plausible explanation for why tumours receiving apparently adequate chemoradiotherapy can nevertheless fail to regress. We acknowledge that the relationship between plasticity and DNA-damage repair is unlikely to be explained by a single pathway. Epigenetic regulation is known to influence both cell-fate plasticity and the DNA-damage response [43, 44, 45], and the marked reduction of 2meH3 [K4] in revCSC may suggest chromatin involvement. At the same time, our single-cell PTM data indicate that signalling processes such as cell-cycle entry, checkpoint activation, Hippo signalling, and stress signalling are closely coupled to stem cell identity and transdetermination. The mechanism linking revCSC identity to DNA-damage resolution is therefore likely to involve a dynamic interaction between signalling state, chromatin state, and DNA-repair pathway choice. Defining how revCSC rapidly resolve therapy-induced DNA-damage is now an important question for the field.

Our findings also suggest that CRT does not merely select pre-existing resistant cells, but can rapidly remodel the epithelial stem cell admixture. We find that FOLFOX, FOLFIRI, and irradiation can polarise rectal cancer epithelia towards a revCSC-dominant state. This shift is not simply explained by apoptosis of proCSC, suggesting that CRT can induce proCSC to revCSC transdetermination - similar to stem cell plasticity observed during intestinal wound healing and infection [46, 47]. We observe the same directionality in LARC patient biopsies, where nCRT is associated with a rapid reduction in proCSC signatures and a gradual increase in revCSC signatures. Moreover, a high revCSC signature is associated with worse overall survival in LARC. These data suggest that CRT can create the very cell-fate ecology that permits residual disease to persist. In this sense, revCSC are not simply a pre-treatment biomarker of poor response; they are also a treatment-induced persister attractor state.

The tumour microenvironment provides a second route into the same resistant state. CAFs are well known to protect cancer cells from therapy, but the relevant epithelial mechanism is often unclear. Here, we show that CAF-induced chemo-radioprotection is strongly associated with CAF-induced revCSC. CAFs can convert proCSC-dominant, chemo-radiosensitive PDOs into revCSC-rich, chemo-radioresistant admixtures, and this effect is additive with CRT-induced plasticity. This provides a direct mechanistic link between stromal signalling, epithelial plasticity, and CRT resistance. Rather than acting only through generic survival factors, CAFs appear to protect LARC epithelia by increasing access to a repair-competent revCSC state. We recently demonstrated that stromal-induced revCSC is driven by CAF-secreted Prostaglandin E2 (PGE_2_) [22]. This may help explain why stromal-rich rectal cancers respond poorly to radiotherapy and have worse clinical outcomes [36, 38, 39]. More broadly, our data suggest that CRT resistance can be generated cell-autonomously by therapy-induced plasticity, or non-cell-autonomously by stromal induction of the same epithelial fate.

These observations have important therapeutic implications. If proCSC are CRT-sensitive and revCSC are CRT-resistant, then one rational strategy is not simply to intensify genotoxic therapy, but to control the cell-fate context in which genotoxic therapy is received. We tested two conceptually opposite strategies: increasing access to CRT-sensitive proCSC through PI3K activation, and blocking access to CRT-resistant revCSC through YAP/TEAD inhibition. PI3K activation is a deliberately provocative strategy, because PI3K signalling is usually viewed as oncogenic and pro-growth. We found that pulsatile PI3K activation could enrich proCSC and increase radiosensitivity in a subset of PDOs. This supports the idea that enriching a chemoradiosensitive cell-fate can create a short-term therapeutic vulnerability. Signalling hyperactivation is an exciting new therapeutic modality and understanding how to deploy signalling activators to treat cancer is an emerging research area [48]. Our results suggest that signalling hyperactivation has the potential to improve CRT responses, but further research is required to identify which patients and cells are most vulnerable, develop optimal timing strategies when combined with CRT, and understand which other pathways provide therapeutic signalling activation opportunities (e.g. WNT/β-Catenin). By contrast, YAP/TEAD inhibition provided a more generalisable strategy to block access to the persister features of the revCSC state. The TEAD palmitoylation inhibitor VT107 (Vivace Therapeutics) [28] increased IR-induced apoptosis across most PDOs, reduced IR-induced access to revCSC, resensitised CAF-protected co-cultures, and could limit access to chemotherapy-induced DNA-repair states. Together, these data suggest that therapies to control cell-fate can convert CRT-induced DNA-damage from a tolerated insult into a lethal one.

LARC and CRC are often conceptualised as WNT signalling-driven diseases, where cell-autonomous APC-mutations prevent degradation of β-Catenin to support tumour proliferation [49]. However, our data indicate that CRT-resistant cells are not the most proliferative cells, but the cells able to retreat into a regenerative revCSC programme capable of withstanding extreme DNA-damage. This creates an important therapeutic distinction: targeting tumour growth and targeting treatment persistence are discrete problems. proCSC drive expansion, but they are vulnerable to CRT. revCSC are less proliferative, but they are better adapted to survive genotoxic stress. Once established, revCSC are difficult to target with standard-of-care therapies [42]. However, when revCSC are induced by CRT or CAFs, blocking plasticity before or during this transition can maintain cells in a more CRT-sensitive state. This provides a therapeutic rationale for combining conventional CRT with agents that either eliminate revCSC-like cells directly (e.g. cellular therapies [33] or antibody-drug conjugates [50]) or restrict proCSC to revCSC transdetermination.

The strongest example of this principle in our study is YAP/TEAD inhibition. Our data reveal that revCSC survive chemoradiotherapy not by avoiding DNA-damage, but by repairing it: isogenic proCSC and revCSC sustain equivalent pH2AX^+^ double-strand breaks, yet only proCSC translate this damage into apoptosis. We therefore speculate that the revCSC state provides a YAP-dependent, repair-competent refuge into which LARC cells retreat under treatment, and that VT107 sensitises cells to CRT by blocking access to this refuge. This interpretation is consistent with the close coupling between Hippo signalling and the DNA-damage response: TEAD transcription factors associate directly with DNA-repair proteins to facilitate recovery from genotoxic stress [51], YAP/TEAD inhibition radiosensitises other tumours by disabling DNA repair and survival signalling [52], and loss of YAP/TAZ is sufficient to convert DNA-damage into cell death [53]. Conversely, activation of YAP is sufficient to protect tumours from therapy-induced DNA damage [54]. Notably, YAP/TAZ are also required for epithelial regeneration following intestinal irradiation [55], the same regenerative programme that revCSC appear to hijack to survive CRT. Our own data reinforce this model: VT107 is selectively toxic to PDOs with high baseline DNA-damage, elevates IR-induced pCHK1 activation, reduces therapy-induced access to revCSC, and collapses the cycling DNA-repair state that chemotherapy otherwise sustains in stromal tumour models.

We introduce SPARTA analysis of patient-derived assembloids to convert spatial transcriptomics from a low-throughput descriptive method into a screening-scale perturbation platform. SPARTA allowed us to measure single-cell chemotherapy responses in stromal human tumour models and revealed that FOLFOX and FOLFIRI do not simply eliminate cycling cells. Instead, chemotherapy-treated stromal assembloids contained epithelial cells with features of both cell-cycle activity and DNA-repair, consistent with a slow-cycling drug-tolerant persister state. While VT107 alone had little effect on established stromal assembloids, YAP/TEAD inhibition blocked access to the actively cycling DNA-repair state and redirected epithelia towards growth arrest and stress. These data suggest that YAP/TEAD activity is required for the drug-tolerant properties of revCSC, and demonstrate that the same cell-fate logic observed in PDOs can be recovered in spatially organised human assembloids.

In summary, our data support a model in which LARC response to CRT is determined by whether genotoxic damage is imposed on a proCSC state that dies, or a revCSC state that repairs and persists. CRT and CAFs both increase access to revCSC, thereby creating a repair-competent refuge that can survive standard-of-care therapy. Pharmacological rewiring of cell-fate, either by increasing access to proCSC or blocking access to revCSC, can restrict this escape route and convert CRT-induced DNA-damage into apoptosis. These results suggest that combining conventional CRT with cell-fate-control therapies may provide a rational route to sensitise poor-response rectal cancers.

### Limitations of the Study

First, although PDOs, PDO-CAF co-cultures, and PDAs capture key epithelial and stromal features of LARC, they do not model the full immune, vascular, microbial, and anatomical complexity of rectal tumours *in vivo*. Second, our irradiation and chemotherapy schedules are designed for systematic perturbation and mechanistic resolution, and do not fully reproduce the temporal structure of clinical fractionated radiotherapy or total neoadjuvant therapy. Third, while measuring pH2AX, pCHK1, pCHK2, and pP53 at single-cell resolution in thousands of organoid cultures provides a comprehensive assessment of DNA-damage response, the precise DNA-repair pathways used by proCSC/revCSC remain an outstanding question for the field. Fourth, drug dose timings were applied uniformly to enable direct comparisons across the cohort. Signalling is a dynamic process and drug-specific temporal dosing strategies are likely to improve efficacy. Finally, YAP/TEAD inhibition requires careful *in vivo* testing, particularly because YAP/TAZ also contribute to normal intestinal regeneration after injury. However, the related clinical-stage TEAD inhibitor VT3989 is currently in Phase 1/2 trials for solid tumours [56] and our study provides rationale to test TEAD inhibitors in combination with CRT to treat LARC. Future studies should define the scheduling, therapeutic window, and normal tissue consequences of combining plasticity-targeting agents with CRT.

## Methods

### Organoid Culture

The PDO culture protocol has been adapted from a previous publication [21]. Briefly, PDOs are thawed and expanded initially in 40 µL growth-factor reduced Matrigel (Corning, 354230, Lot number 12924002) in 6-well plates supported by 2.5 mL media. Organoid growth media contained Gibco Advanced DMEM/F-12 (Thermofisher, 12634028) with 25% WRN-conditioned media (WNT-3A, R-Spondin, Noggin), 50 ng/mL recombinant mEGF (PeptroTech, 315-09-100UG), 500 nM A83-01 (Cambridge Bioscience, 909910-43-6), 10 µM SB-202190 (VWR, CAYM10010399), 1% Gibco serum-free N2 100X (Thermofisher, 17502048), 1% Gibco serum-free B27 (100X, Thermofisher, 17504044), 10 mM nicotinamide (Sigma-Aldrich, N3376), 1 mM N-acetyl-L-cysteine (NALC) (Sigma-Aldrich, A9165), 10 mM HEPES (Sigma-Aldrich, H3375), 2% Hyclone penicillin/streptomycin/amphotericin B (Cytiva, SV30079.01), and 1% Gibco Glutamax 100X (Thermofisher, 35050061). ROCK inhibitor Y-27632 (10 µM Sigma-Aldrich, Y0503) was added for 24 hours after thawing and single-cell dissociation. PDOs were passaged once weekly. After at least 2 passages from thawing, PDOs were dissociated using 1 mL Gibco Try-PLE (Thermofisher 12604013) per 6-well plate at 37°C, 5% CO_2_, and expanded using the 5% Matrigel method, whereby single cells were resuspended in 1 mL Matrigel, then diluted in 19 mL media without mEGF or WRN, and cultured in an ultra-low attachment T75 flask (Corning, BC371). Four days after the expansion in 5% Matrigel, the media was changed for ‘Base Media’ containing only Gibco Advanced DMEM/F-12, NALC, Hyclone Pen/Strep/Amphotericin and Gibco Glutamax. On the fifth day of 5% Matrigel culture, PDOs were harvested for experimental seeding in 96-well plates.

### CAF Culture

CAFs were obtained from the De Wever laboratory (University of Gent) [57]. CAFs were thawed and grown in Gibco DMEM (Thermofisher, 11965092) containing 10% Gibco Fetal Bovine Serum (Thermofisher, 10082147) and Hyclone penicillin/streptomycin/amphotericin B (Cytiva, SV30079.01). 3 million CAFs were seeded into a T175 flask in 20 mL media. To passage, CAFs were lifted using 4 mL Gibco TryPLE per flask at 37°C for 1 minute before adding 16 mL media per flask. Lifted CAFs were spun down at 1,100 RPM for 4 minutes before seeding at 3 million cells per T175 flask. In the final expansion phase before seeding, CAFs are passaged and grown in DMEM containing 2% FBS for 3 days, at which point they are ready to be harvested for an experiment.

### PDO-CAF Perturbation Arrays

On day 0, PDOs were harvested from 5% Matrigel culture using ice-cold PBS and seeded in 50µL droplets into 96-well plates at a density of 1500 PDOs per well. In co-culture conditions, CAFs were harvested and mixed with PDOs at 3000 CAFs/ µL, before seeding as above. 300 µL Base Media was added to each well. On day 1, the media was replaced and the following drug treatments were added: 2 µM 5-FU (Sigma-Aldrich, F6627), 100 nM oxaliplatin (Sigma-Aldrich, O9512), 10 nM SN-38 (Sigma-Aldrich, H0165), 5 µM UCL-TRO-1938 (Vanhaesebroeck Laboratory, UCL), 4 µM VT107 (Vivace Therapeutics), and 4 µM Nintedanib (Sigma-Aldrich, SML2848), or DMSO vehicle control. On day 2, PDOs were taken for either sham or 16 Gy X-ray radiation using the CIX3 (XStrahl) irradiator at a dose rate of 3.14 Gy/min, after which Base Media was replaced. Cells were fixed at 3 hours after chemoradiation (day 2), or 48 hours after chemoradiation (day 4). Daily 100% Base Media changes were continued up until the point of fixation.

### Thiol-reactive Organoid Barcoding *in situ* Mass Cytometry (TOB*is* MC)

All cultures were prepared based on the protocol outlined in Sufi and Qin *et al*, Nature Protocols [30]. Prior to fixation, 25 µM 5-iodo-2’deoxyuridine (^127^IdU) (Sigma-Aldrich, I7125) was added to cultures and incubated at 37°C for 25 minutes, after which 1% Protease Inhibitor Cocktail (Sigma-Aldrich, P8340) and 2.5% PhosSTOP (Roche, 4906845001) were added for a further 5 minutes of incubation. Subsequently, cultures were fixed using 200 µL/well 4% paraformaldehyde (Thermo Fisher, AAJ19943K2) and barcoded using 126-plex thiol-reactive barcodes overnight at 4°C. To enable debarcoding of cells treated with platinum based chemotherapies (e.g. FOLFOX), we swapped ^196^Cisplatin and ^198^Cisplatin used in TOB*is* 1.0 [29, 30] for 6 µM mDOTA-^103^Rh and 0.4 µM BABE-^108^Pd (hereafter referred to as TOB*is* 2.0). TOB*is* 2.0 barcodes were quenched with 1 mM glutathione (Sigma-Aldrich, G6529), after which cultures were pooled and dissociated into single-cells using the Gibco 1 mg/mL Dispase II (Thermofisher 17105041), 0.2 mg/mL Gibco collagenase IV (Thermofisher 17104019), and 0.2 mg/mL DNase I (Sigma-Aldrich, DN25) in GentleMACS M-tubes on GentleMACS Octo dissociators. Samples were stained using the antibodies described in Table 2, using 0.1% Triton X-100 to permeabilise the cells before intracellular staining. Stained samples were treated with 0.1% Iridium intercalator (Standard Biotools), washed sequentially in PBS with 0.1% BSA, PBS, and distilled water, then stained with 25 µg/mL bis(2,2’-bipyridine)-4’-methyl-4-carboxybipyridine-ruthenium N-succinimidyl ester-bis(hexafluorophosphate) (Sigma-Aldrich, 96631) per 10 million cells in 0.1 M NaHCO_3_. EQ6 beads (SBT) were added in a ratio of 1:4 on the CyTOF-XT (SBT). Samples were diluted to achieve an acquisition event rate of 400 events / second.

### Single-cell RNA-seq Data Analysis

Single-cell RNA-seq data was acquired as described in Nattress *et al*., [33] and analysed as described in Molyneux and O’Sullivan *et al*., [22]. Putative signalling pathway inference was performed using PROGENy [34].

### PDO Chemotherapy LD_50_ Dose-responses

PDOs were harvested from culture 5 days after passage, mixed thoroughly to ensure even distribution, and seeded into clear-bottomed 96-well plates (Greiner, 655098) in technical triplicates. Cultures treated with 10 doses of 5-FU, oxaliplatin and SN-38 for 48 hours. Lethal (50% DMSO) and vehicle controls were also seeded in triplicate for each drug treatment. After drug treatment, all media was removed and 50µL CellTiter-Glo 3D (Promega, G9681) was used to assess viability according to the manufacturer’s protocol. Luminescence readings were taken using the Varioskan Lux plate reader (Thermofisher). Viability (%) was calculated by subtracting the background luminescence of the lethal control, then dividing by the luminescence of the untreated control. LD_50_ calculations were performed by fitting the % viability to non-linear, variable slope dose-response curve using GraphPad Prism v10.1.1.

### CyTOF Data Analysis

FCS files were uploaded to Cytobank (Beckman Coulter) where gating was performed for Gaussian parameters, DNA (Ir intercalator) and cell type (epithelia: PanCK and EpCAM; CAFs: Vimentin and FAP). Events for each cell type were exported and batch normalisation was performed with CytoBatchNorm [58], using internal controls spiked into each batch as anchors. Marker values were arcsinh transformed, then normalised to cell size [59]. All subsequent analyses, including Trellis/TreEMD [21] and PHATE [60] were performed in JupyterLab v4.3.4.

### Calculation of Stem Cell Index

Candidate proCSC and revCSC markers were selected from existing literature [12, 17] and single-cell RNA-sequencing of 9 colorectal PDOs. Those that passed validation at protein level in TOB*is* MC were used to generate a proCSC score and revCSC score per cell by z-scaling, then taking the mean, of the pertinent markers (proCSC markers: EPHB2, RRM2, CDK1, Ki67, TOP2A; revCSC markers: L1CAM, CD55, CK18, ANXA1, OPTN, TROP2, BIRC3). The stem cell index (SCI) was calculated by subtracting the revCSC score from the proCSC score.

To isolate treatment-induced changes in SCI (ΔSCI), we computed an adjusted ΔSCI to correct for any time-related component of Δ SCI in each PDO as follows:

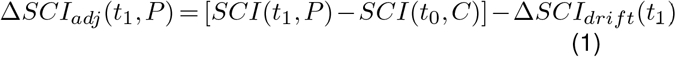

where Δ*SCI*_*adj*_ is the PDO- and perturbation-specific change in SCI over time accounting for any drift in controls over time. *t, P*, and *C* denote time point, perturbation, and control respectively, and Δ*SCI*_*drift*_ is given by:

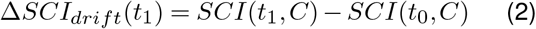

### Conditional Flow Match Treatment Effect Modelling with GeoSinkhorn Flow

Geodesic Sinkhorn [32] is a graph-based optimal transport (OT) method that implements the fast Sinkhorn algorithm for optimal transport on graphs using a heat-geodesic kernel. GeoSinkhorn Flow is a neural network variant of Geodesic Sinkhorn that not only generalises OT computations to new distributions not seen during training, but also uses flow matching to recover continuous transport paths. GeoSinkhorn Flow employs a heat-geodesic embedding [61], which provides a heat-geodesic distance-preserving Euclidean embedding of the data. It then performs Euclidean flow matching in this latent space. For more details, see Xu, Wilkinson, Ni *et al*. (in preparation).

GeoSinkhorn Flow additionally supports the use of conditional treatment signals to predict flows from control to treatment and to generalise across treatments. In this manuscript, we applied GeoSinkhorn Flow to compute optimal transport paths between proCSC- and revCSC-gated populations for the 16 Gy and 16 Gy + FOLFOX conditions across 10 PDOs. From the resulting flows, we show changes in the expression of IdU, pH2AX [S139], and cPARP [D214] relative to the DMSO control.

The flow-matching component of GeoSinkhorn Flow uses pseudotime bins and constructs a transport path that passes through all bins sequentially at the distribution level. This approach provides more accurate transport paths and higher temporal resolution than using only two time points. Therefore, rather than treating the control and perturbation conditions as two pseudotime bins, we introduce additional intermediate bins using the MELD method [62].

MELD compares single-cell data from control and treatment samples by calculating, for each cell, the relative likelihood it would be observed under each condition. It builds a graph connecting similar cells and smooths the condition labels across it, via graph signal processing, to estimate this likelihood, revealing which cells are most enriched or depleted by the treatment at single-cell resolution. Pseudo-time bins are constructed to span across the MELD score gradient so that the trajectories are trained in alignment with the treatment effect.

### S:CORT Data Access and Clinical Information

Details of the S:CORT cohort and consortium have been reported previously [63]. S:CORT clinical data and raw gene counts were obtained following an approved request to the S:CORT data access committee. The counts comprised probe-level counts summarised at the gene level. After mapping, probes without an assigned gene symbol were excluded. Where multiple probes mapped to the same gene, probes were collapsed to a single gene-level value by retaining the probe with the highest mean expression across all samples. The resulting gene-level expression matrix was then used for downstream signature scoring. Colon cancer cases were filtered out and 223 rectal cancer patients who underwent neoadjuvant chemoradiotherapy were retained.

### Standard-of-Care Glasgow Serial Sampling Cohort and Ethics

Locally advanced rectal cancer patients with T stage 2-4 were recruited for a serial sampling biopsy collection study during their standard of care neoadjuvant radiotherapy or chemoradiotherapy. Patients over 18 years of age and scheduled for radiotherapy at the Beatson West of Scotland Cancer Centre or Lanarkshire hospitals were enrolled. Patients were excluded if they had bleeding conditions, anti-coagulant prescriptions, or tumours at or below the dentate line. Ethics approval was granted by NHS Glasgow and Greater Clyde research ethics committee (ref: 18WS0003). The study was funded by Varian Medical Systems, Inc., A Siemens Healthineers Company, and the Beatson Cancer Charity.

Patients were stratified into the following neoadjuvant regimens: short course radiotherapy (5 × 5Gy) with or without chemotherapy; or long course radiotherapy (25 × 1.8-2.1Gy) with or without chemotherapy. Chemotherapy consisted of either capecitabine, capecitabine and oxaliplatin (XELOX), 5-FU and folinic acid, or folinic acid, 5-FU, and oxaliplatin (FOLFOX), given between weeks 2-4 of their neoadjuvant therapy. Biopsy samples from a total of 48 patients were collected at four time points: week 0 before radiotherapy (*n* = 34), and at weeks 2 (*n* = 31), 6 (*n* = 26), and 12 (*n* = 29) after commencement of neoadjuvant treatment. Areas sampled were macroscopically identified as malignant epithelium. Tissues were processed into formalin-fixed paraffin-embedded (FFPE) tissue slides at the NHS Greater Glasgow and Greater Clyde Biorepository at the Queen Elizabeth University Hospital.

### RNA Extraction, Library Preparation, and RNA-Sequencing

RNA sequencing was conducted using 10 µM thick FFPE tissue sections, which were submitted for sequencing at GENEWIZ (Azenta Life Sciences, Leipzig, Germany). RNA extraction was performed with the RNeasy FFPE kit (Qiagen, 73504) and RNA quality was evaluated using an Agilent Bioanalyzer RNA 6000 Nano Chip. Ribosomal RNA depletion was carried out using the NEBNext rRNA Depletion Kit v2, followed by library construction with the NEBNext Ultra II RNA Library Preparation Kit. Paired-end sequencing (2 × 150 bp) was performed on an Illumina NovaSeq platform, generating raw FASTQ files.

FASTQ reads were assessed and filtered for quality using FastQC (v0.12.1), MultiQC (v1.17), and fastp (v0.23.4). The processed reads were then pseudo-aligned to the GRCh38 reference transcriptome and quantified using Kallisto (v0.50.1) to yield gene level counts. All samples achieved more than 2 million pseudo-aligned reads, a duplication rate below 50%, and showed no outliers on PCA, and were therefore used for down-stream analysis. Samples from 48 patients were analysed, including 34 samples for week 0, 31 for week 2, 26 for week 6, and 29 for week 12, and 16 matched samples across all time points.

### ssGSEA

Single-sample gene set enrichment analysis (ssGSEA) was performed using the GSVA package (version 2.0.7). For the standard-of-care cohort, variance-stabilised transformed gene expression values were generated using DESeq2 (1.46.0) and used as input. For the S:CORT cohort, the processed mean gene-level expression matrix was used as input. Genes in the proCSC and revCSC signatures were intersected with those present in the corresponding expression matrix, and ssGSEA enrichment scores were calculated for each sample using ssgsea-Param with alpha = 0.25, minSize = 1, maxSize = Inf, and normalisation enabled. Paired Wilcoxon signed-rank tests were used to assess changes between time points, and significance was defined using p< 0.05.

### Survival

The ssGSEA enrichment scores for proCSC and revCSC gene sets were used with patient overall survival and disease-free survival data to determine their prognostic value. The “surv_cutpoint” function from the package “survminer”(version 0.5.2) was used to dichotomise the cohort into high and low enrichment groups based on ssGSEA scores for proCSC or revCSC separately. The “minprop” parameter was set to 0.15 to ensure meaningful group sizes, and the function automatically calculated an optimal threshold. The functions “Surv” and “survfit” from the package “survival” (version 3.2.13) were used to fit a Kaplan Meier curve to the dichotomised dataset with either overall or disease-free survival, where death or recurrence indicated an event respectively. The function “ggsurvplot” from survminer was used to plot the curves with 95% confidence intervals. The “coxph” function from the survival package was used to apply a univariate Cox proportional hazards regression model on the dichotomised data; p value and hazard ratio are reported on the plot.

### Patient-Derived Assembloid (PDA) Establishment and Culture

Technical aspects of generation of murine intestinal assembloids are described by Lin *et al*., [40], to which we applied minor adaptations to establish human patient-derived assembloids. Briefly, PDOs were harvested from 40 µL Matrigel droplets 4 days after passaging using ice-cold PBS. CAFs were harvested and split into 2 halves. PDOs were checked for density by manual counting, added to one half of the CAFs. Both samples were then resuspended in Matrigel, ensuring a final concentration of 40,000 CAFs/µL in the CAF-Matrigel mix and an equal concentration of CAFs plus 2-3 PDOs/µL in the CAF-PDO-Matrigel mix. Sterile 90 mm Petri dishes (Thermo Scientific, 11759252) were lined with Parafilm M (Merck, HS234526B) and left to air-dry under a sterile hood before being placed over ice. 2 µL CAF-Matrigel droplets were initially pipetted onto the Parafilm-lined petri dish over ice, followed by quick but gentle pipetting of a 2 µL CAF-PDO-Matrigel droplet into the centre of the CAF-Matrigel droplet. Droplets were covered and incubated at 37 °C for 10 minutes, then individually transferred with care to a 12-well plate containing 1.5 mL Base Media per well. 10 assembloids were transferred per well. Assembloids were subsequently grown in Base Media at 37 °C, 5% CO_2_ on an orbital shaker (Edmund Buehler, Miniature Shaker KM CO2-FL) at 170 RPM with daily 100% media changes.

### Spatial Perturbation of ARrayed Tumour Assembloids (SPARTA)

Assembloids were subject to drug treatments on day 4 after seeding at the same doses as the co-culture perturbation arrays. Drugs were removed on day 5, and assembloids were fixed in 4% PFA 1 mL/well on day 6 for 2 hours at 21°C. Fixed specimens were processed using a formalin-fixed paraffin-embedded (FFPE) protocol optimised for small tissues. Briefly, samples were dehydrated through a graded alcohol series (5 h 30 min), cleared in xylene (3 h), infiltrated with paraffin (4 h), and embedded in Type H Paraffin Wax (Epredia, 8338). Each treatment group was embedded as a separate FFPE block. Tissue microarrays (TMAs) were constructed using the automated TMA Grand Master (3DHISTECH) system according to the manufacturer’s instructions. Five cores (1.0 mm diameter) were sampled from each donor assembloid FFPE block and transferred to a recipient block containing Type H Paraffin Wax (Epredia, 8338). Cores were arranged in a 5×12 array with 0.5 mm spacing for compatibility with the capture area of Xenium slides (10X Genomics), with one blank to create a unique orientation. Following TMA construction, blocks were incubated overnight at 37 °C and annealed by four cycles of alternating 1 h incubations at 4 °C and 37 °C [64]. Annealed TMA blocks were sectioned at 5 µm using a rotary microtome and mounted onto Xenium slides. Tissue sections and slides were prepared, dried and stored according to the manufacturer’s recommendations prior to Xenium processing. Slides were subsequently processed on Xenium Analyzer (10X Genomics) using the 5K Xenium Prime pan-tissue assay kit. Following Xenium analysis, slides were stained with haematoxylin and eosin to reveal tissue architecture.

Data was analysed using Scanpy v1.11.5 [65] and Squidpy v1.6.5 [66] on JupyterLab v.4.3.4. QC cutoffs of total transcripts < 100 and total genes <50 were used. Transcriptional signatures were curated from literature, and also from Over-Representation Analysis using Gene Ontology Biological Process Annotations 2025 [67]. Assembloids were visualised using Xenium Explorer (gene signatures can be found in Table S1). Differentially expressed genes in each Leiden cluster can be found in Table S2, and signatures curated from Gene Ontology can be found in Table S3.

### Statistical Analyses

All statistical analyses were performed using Graph-Pad Prism 10.1.1, or the scipy.stats package v1.18.0 on JupyterLab v.4.3.4. P-values of less than 0.05 were considered statistically significant.

## Supporting information

Table S1 (Xenium Gene Groups)

Table S2 (DEGs per Leiden Cluster)

Table S3 (Signatures Derived from Gene Ontology ORA)

## Data Availability

Raw and processed CyTOF data and illustrations are available as a Community Cytobank project 127111 (https://community.cytobank.org/cytobank/experiments/127111).

Time course specific data for Figure 3 are available as a separate Community Cytobank project 127114

(https://community.cytobank.org/cytobank/experiments/127114). Processed Xenium data is available at Zenodo (https://zenodo.org/records/21103087).

## Acknowledgements

We are extremely grateful to M. Garnett, H. Francies and the Cell Model Network UK for sharing CRC PDOs and Oliver De Wever for sharing CRC CAFs. We thank Y. Guo and the UCL CI Flow-Core for CyTOF support. We acknowledge the Translational Technology Organoid Platform for deriving and supplying the organoid models to use in this research and the UCLH Biobank for Studying Health and Disease for providing the human tissue samples and the clinical data that was used in and supported the generation of the organoid models. Transcriptional data from patient samples was generated by the stratification in colorectal cancer consortium (S:CORT) consortium (consortium membership detailed here: https://scort.org.uk/about-scort.html). Access to this data was facilitated by Dr. Philip Dunne, and was provided via an academic subproposal application. We also acknowledge the Pathology Translational Technology Platform for histology services (RRID:SCR028430), and the Genomics Translational Technology Platform and Imran Uddin for preparing and analysing Xenium slides. We thank Vivace Therapeutics for providing access to VT107. This work was supported by the Cancer Research UK City of London Centre (CTRQQR-2021/100004), Cancer Research UK (C60693/A23783, DRCPFA-May23/100003), UKRI Medical Research Council (MR/T028270/1, APP51437), UCLH Biomedical Research Centre (BRC422, BRC1107), and CRUK CRC-STARS Strategic Grant (SEBCRCS-2024/100001). S:CORT was funded by a UKRI Medical Research Council Stratified Medicine Consortium programme grant (MR/M016587/1) and co-funded by Cancer Research UK. Research in the B.V. laboratory was supported by Cancer Research UK (DRCRPG-Nov24/100006, TICCPP-2022/100005 and DRCTDPJT-Nov21/100001). The Standard of Care Rectal Cancer cohort was funded by Varian Medical Systems, Inc., A Siemens Healthineers Company, and the Beatson Cancer Charity.

## Glasgow Serial Sampling Consortium

Fiza Ishaqwala, Thomas A. Wright, Ashley K. McCulloch, Liang Tang, Chia Y. Kong, Lily V. S. Hillson, Kathryn Pennel, Lynsey Devlin, Ross K. McMahon, Sean M. O’Cathail, Joanne Edwards, Campbell S. D. Roxburgh.

## Author Contributions

N.L. conceived the project, designed the study, made TOB*is* 2.0 barcodes, performed all rectal PDO and assembloid experiments and analyses, and wrote the paper.

F.I. and T.W. performed bulk-RNA sequencing and survival analyses on clinical data.

A.W. performed computational flow matching analysis.

P.V., E.B., and S.V. isolated rectal cancer PDOs and provided organoid culture support.

K.T. developed assembloid array fabrication for SPARTA. R.O’S. acquired and analysed all scRNA-seq data.

S.C. conjugated metal-tagged antibodies.

A.K.M. collected and processed Glasgow tissue samples.

A.D. supported organoid culture and analysis.

S.K. supervised computational flow matching analysis.

B.V. provided reagents and supervised the use of PI3K activation strategies.

Glasgow Serial Sampling Consortium provided longitudinal RNA-seq data of LARC patients treated with nCRT.

C.S.D.R. supervised bulk-RNA sequencing and survival analyses on clinical data.

M.H. supervised chemoradiotherapy treatments and analysis.

C.J.T. conceived and supervised the project, designed the study, analysed the data, and wrote the paper.

## Supplementary Information

### Supplementary Figures

**Figure S1.**
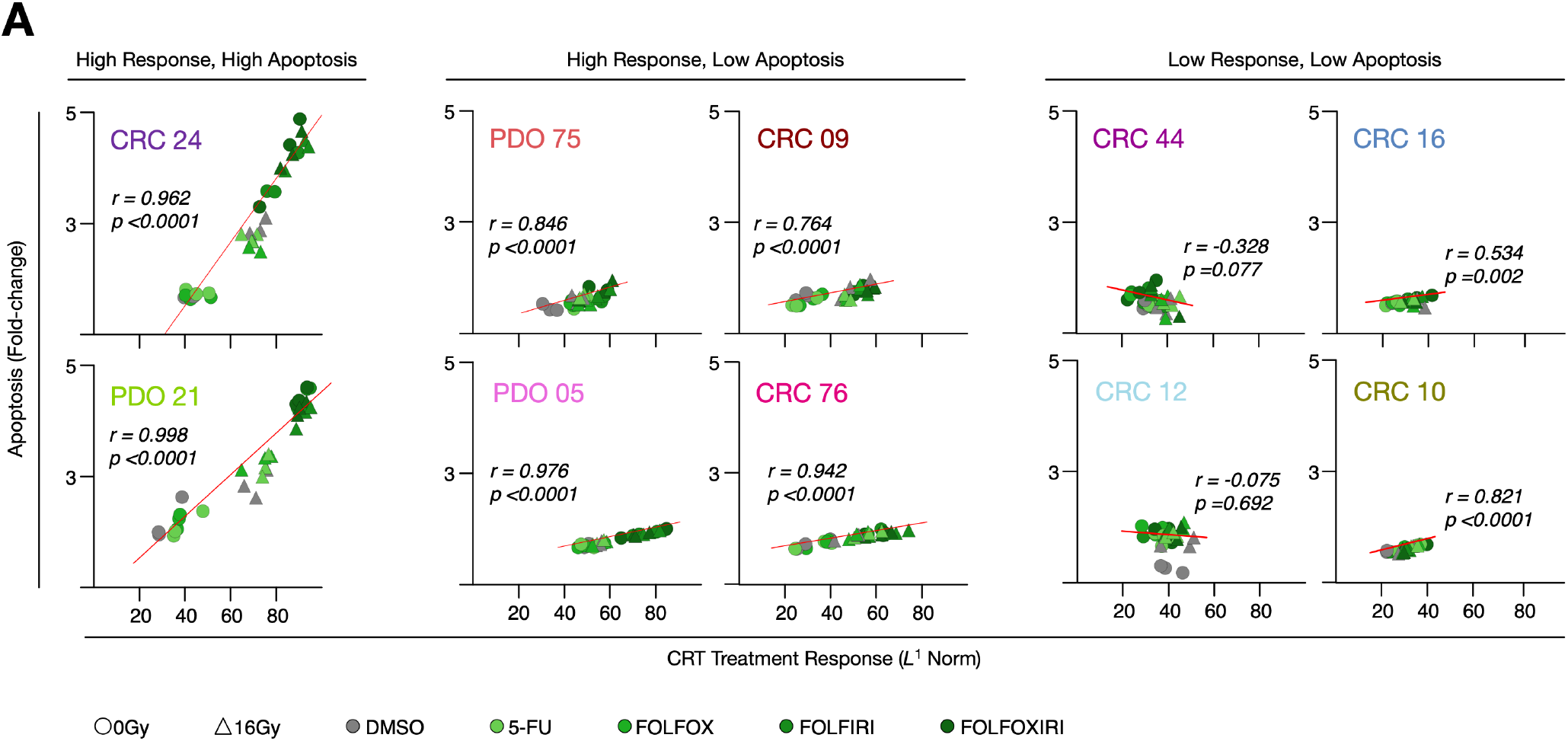
Treatment Responses of Rectal Cancer PDOs to CRT. **A)** Absolute distance between treatment conditions in 10 PDO monocultures captured using the L^1^norm of the Trellis node matrix relative to fold-change apoptosis. Regression lines, Pearson’s coefficient of correlation (r), and p-values were calculated using scipy.stats.linregress (v1.18.0).

**Figure S2.**
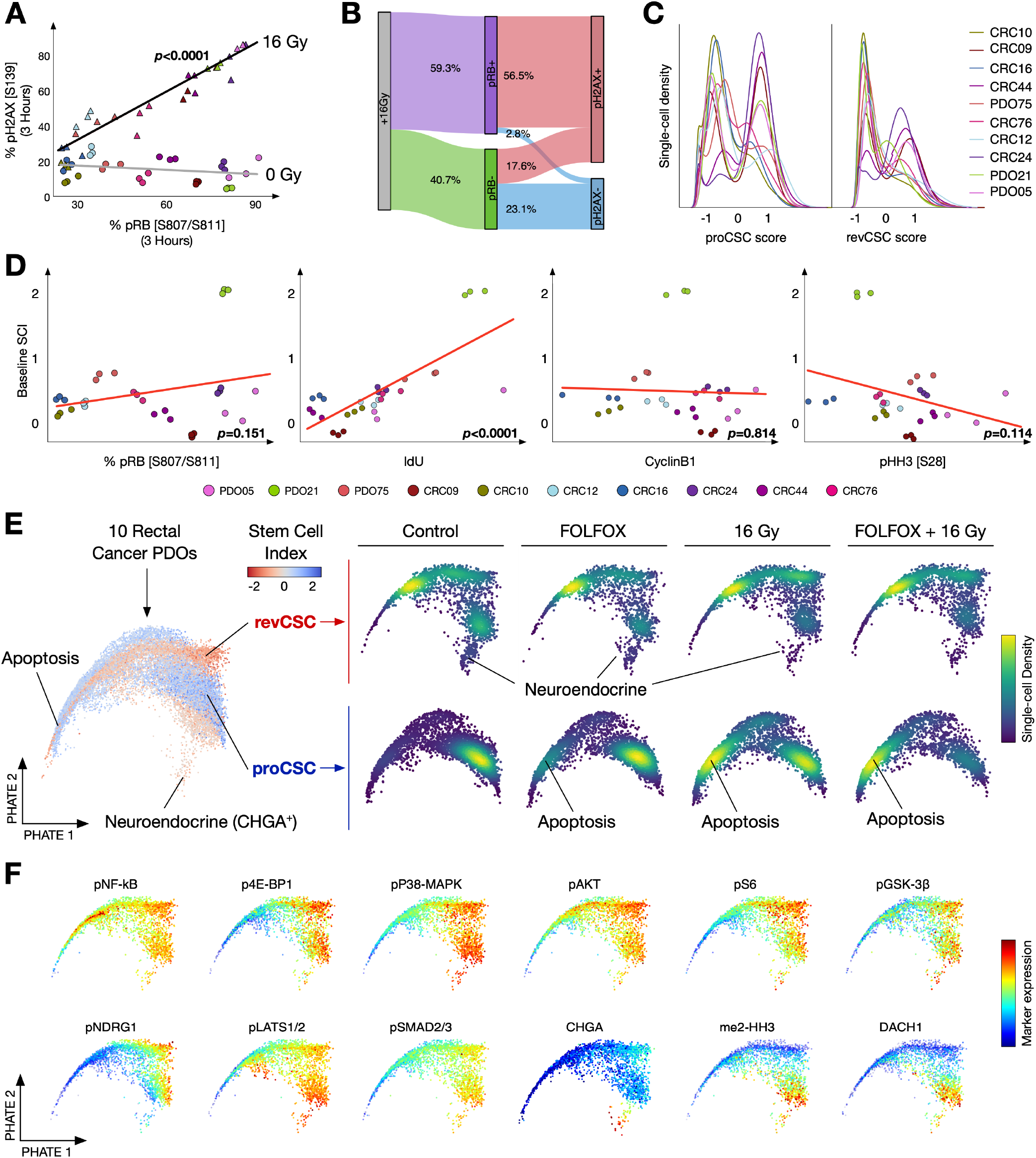
Single-cell Analysis of proCSC- and revCSC-specific CRT Responses. **A)** DNA damage (pH2AX [S139]) against the proportion of actively cycling cells in 10 PDOs treated with +/-IR. **B)** Sankey diagram of DNA-damage by cell-cycle activity in all 10 PDOs. **C)** Single-cell density histograms of proCSC and revCSC scores. **D)** Stem cell index (SCI) against cell-cycle markers. **E)** Single-cell PHATE embeddings of proCSC and revCSC populations from all 10 PDOs, treated with +/-FOLFOX, +/-IR, +/-FOLFOX + IR, coloured by stem cell index (left) or single-cell density per condition (right). **F)** Single-cell PHATE embeddings of signalling and cell-fate markers from 10 PDOs. Regression lines and p-values were calculated using scipy.stats.linregress (v1.18.0).

**Figure S3.**
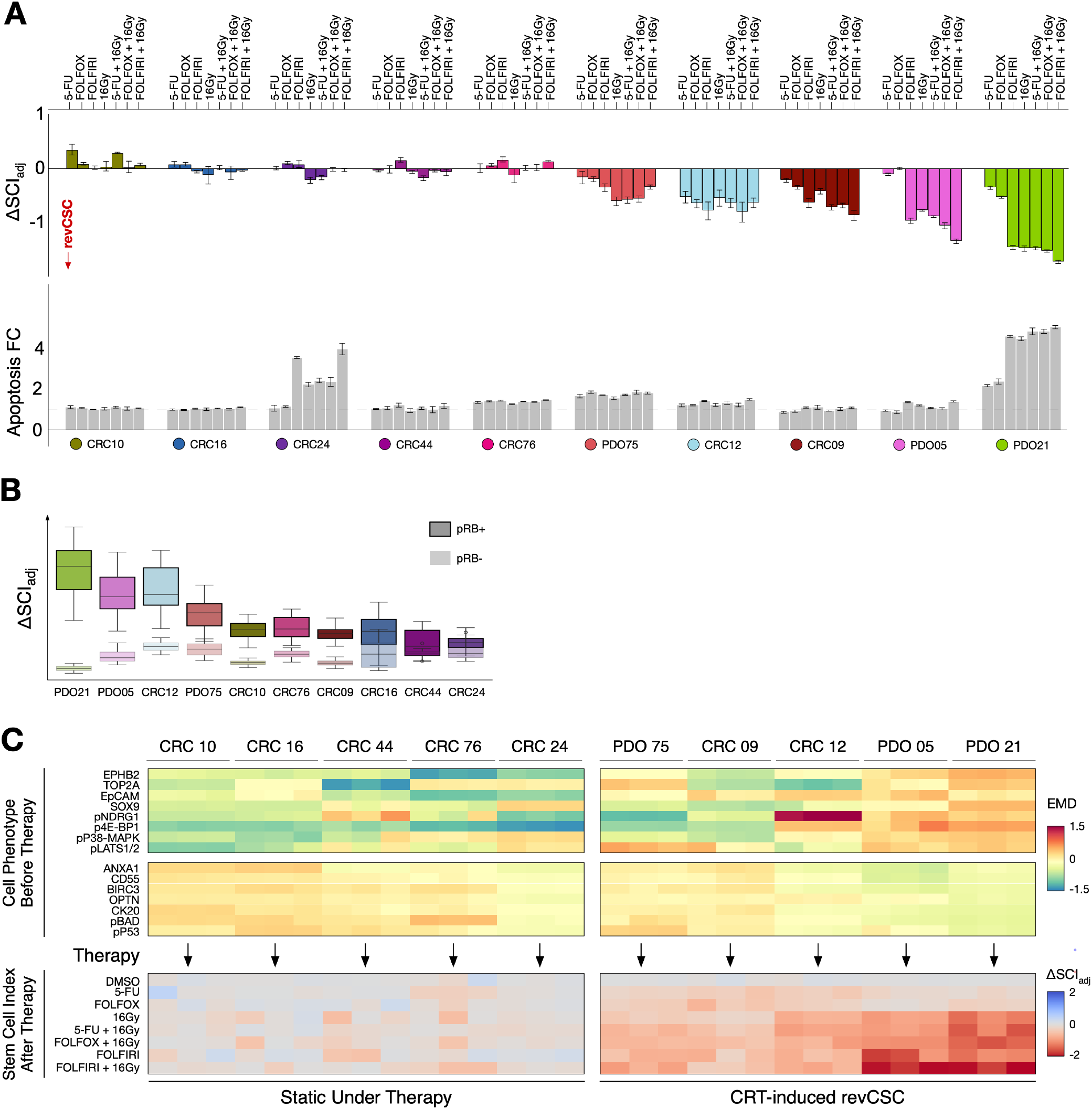
Stem Cell Dynamics Following CRT. **A)** Change in SCI and apoptosis induced by 5-FU, FOLFOX, and FOLFIRI, +/-IR. **B)** CRT-induced SCI changes (absolute values) in pRB^+^ and pRB^-^ LARC cells. **C)** Baseline EMD before treatment (top) and change in SCI after treatment (bottom) for 10 rectal cancer PDOs.

**Figure S4.**
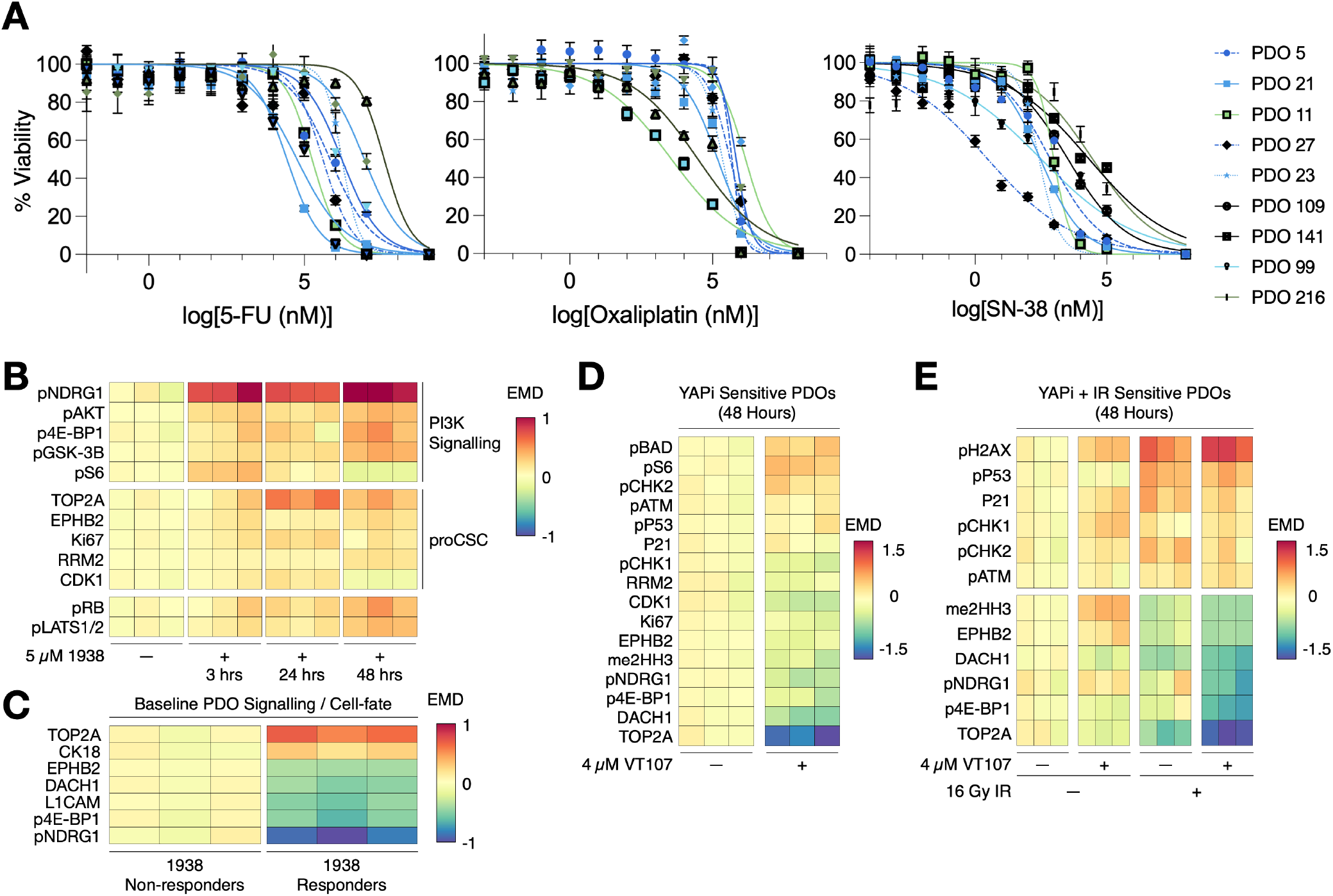
Rectal Cancer PDO Responses to CRT and Signal Rewiring Drugs. **A)** Viability dose-response curves of CRC PDOs to 5-FU, Oxaliplatin, and SN-38, used the calculate the LD_50_ values in Figure 4. **B)** EMD heatmap of PDO21 treated with 5 µM UCL-TRO-1938 and analysed 3, 24, and 48 hours. **C)** EMD heatmap of baseline signalling from grouped 1938 non-responders (CRC44, CRC9) and 1938 responders (PDO05, CRC16, CRC76) PDOs. **D)** EMD heatmap of VT107-sensitive PDOs (CRC24, CRC12, PDO05, PDO75, CRC09) +/-4 µM VT107 (48 hours). **E)** EMD heatmap of VT107-insensitive, but VT107 + IR sensitive PDOs (PDO21, CRC16) +/-VT107 +/-IR.

**Figure S5.**
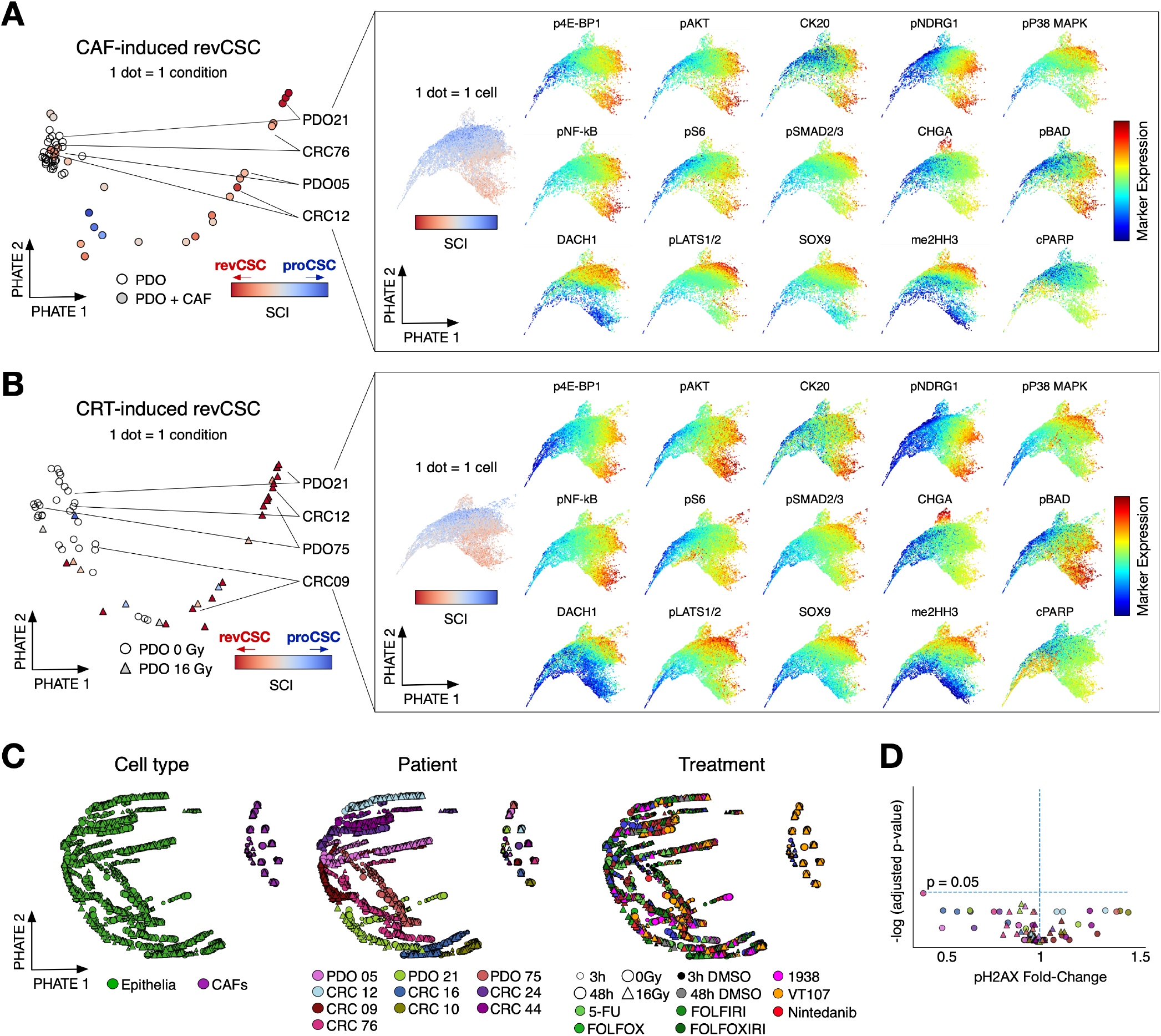
Comparison of CAF-induced revCSC and CRT-induced revCSC. **A)** Trellis-PHATE embedding of PDO and PDO+CAF cultures (epithelial cells only) coloured by stem cell index (left); single-cell PHATE embeddings of CAF-responsive PDOs coloured by SCI and marker expression (right). **B)** Trellis-PHATE embedding of PDO monocultures +/-IR (left); single-cell PHATE embeddings of IR-responsive PDOs coloured by SCI and marker expression (right). **C)** Trellis-PHATE embeddings of PDOs and CAFs across 2,400 cultures coloured by cell type, patient, and treatment. **D)** Volcano plot of relative pH2AX [S139] expression in PDO+CAF cultures +/-IR vs paired PDO monoculture conditions (epithelial cells only). Statistical significance was assessed using multiple paired t-tests with false discovery rate correction for multiple comparisons (*n*=3).

**Figure S6.**
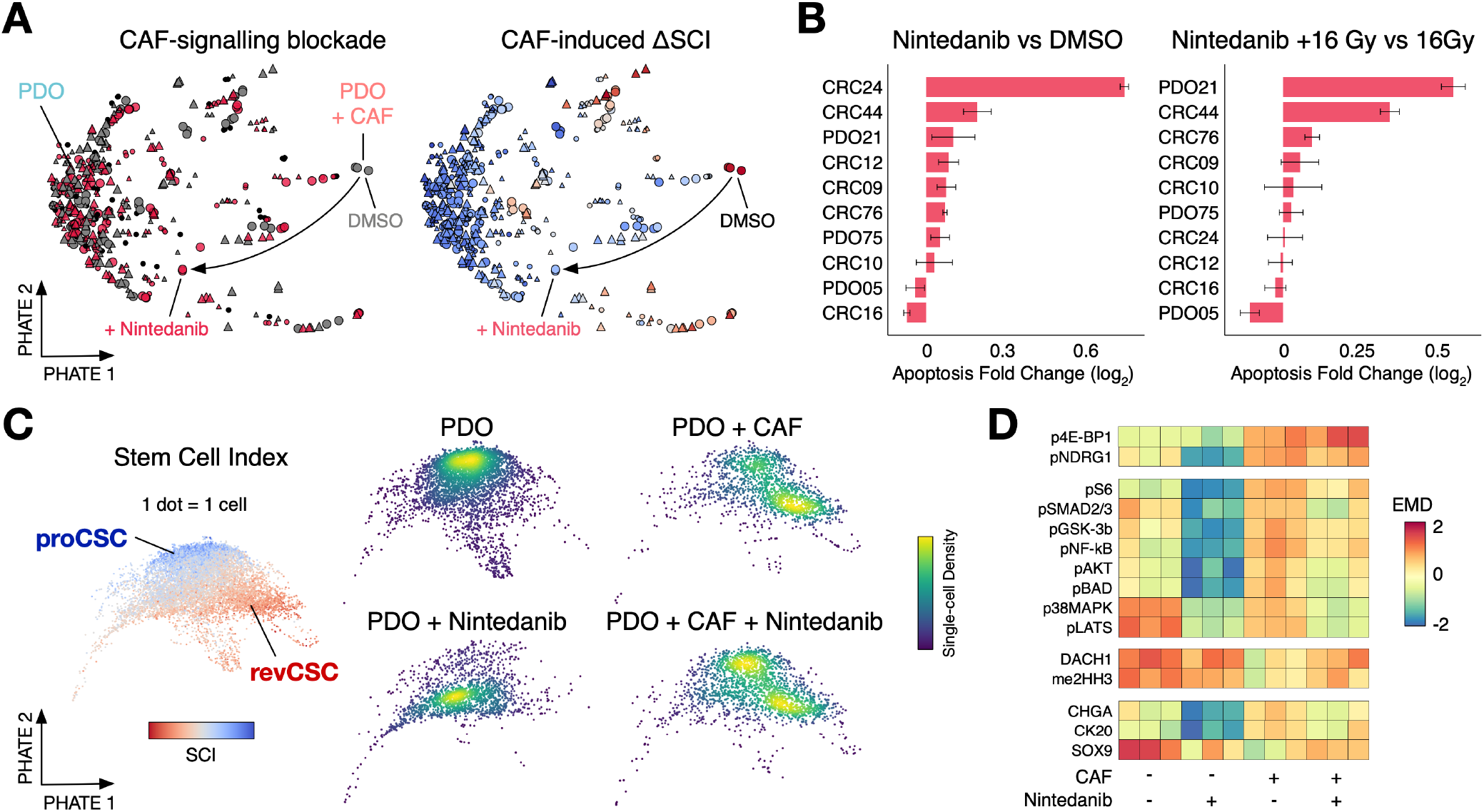
Targeting Stromal Signalling May Radiosensitise LARC PDOs. **A)** Paired TreEMD-PHATE highlighting DMSO (grey) and Nintedanib (red) treated cultures. **B)** Epithelial apoptosis fold-change of Nintedanib-treated PDO+CAF cultures. Error bars denote SEM (n=3). **C)** Single-cell PHATE embedding of PDO21 treated with +/-CAF, +/-Nintedanib. **D)** EMD heatmap of signalling pathways affected by treatment with +/-CAF, +/-Nintedanib across 10 PDOs.

**Figure S7.**
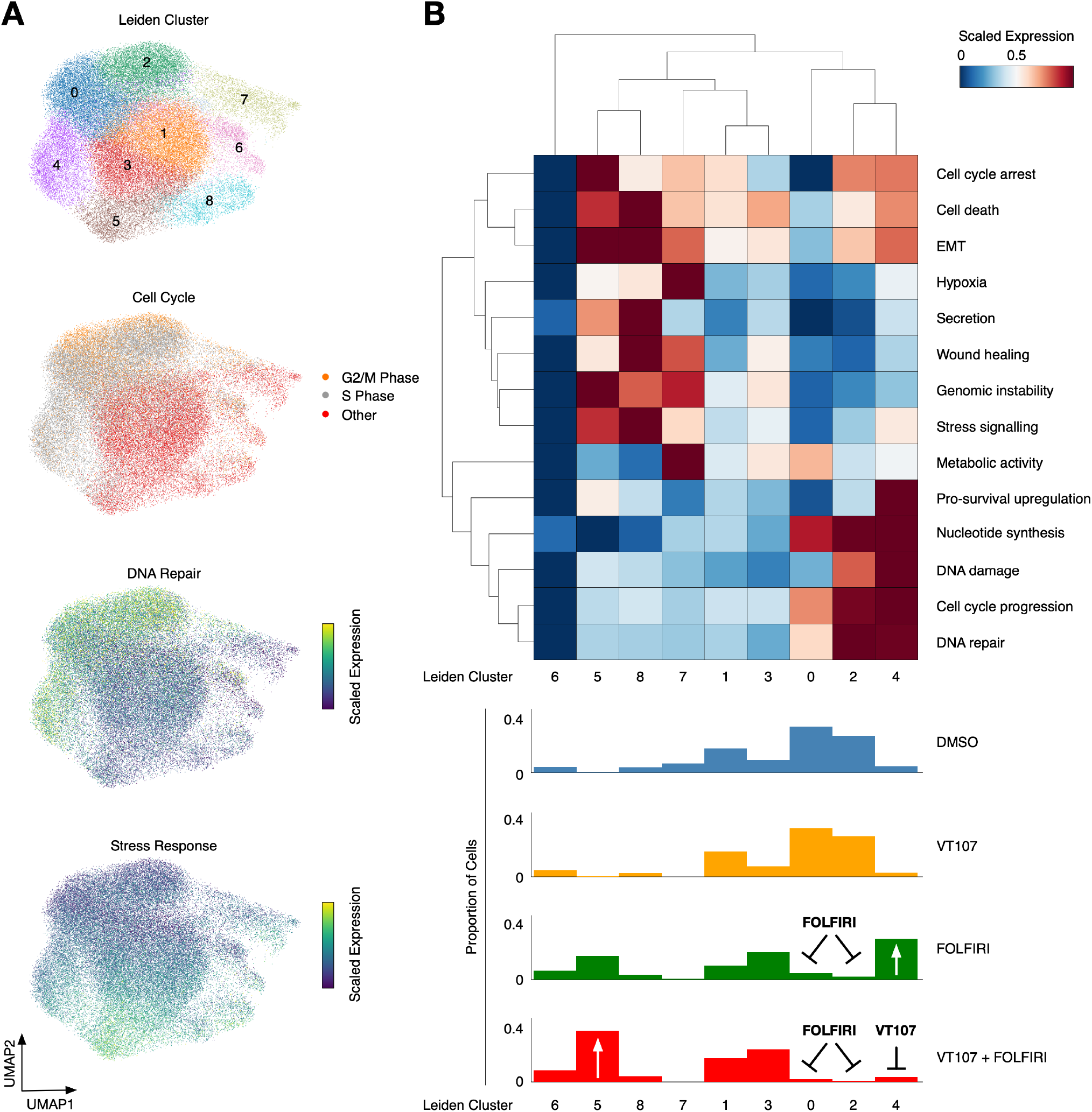
YAP/TEAD Inhibition Attenuates CRT-induced DNA-damage Repair. **A)** UMAP embeddings of single-cell epithelial cell gene expression from SPARTA analysis coloured by Leiden clustering, cell-cycle phase, DNA-damage repair signature, and stress response signature. **B)** Heatmap of transcriptional signatures across Leiden clusters (top) with proportions of cells in each cluster per treatment condition (bottom).

